# Mitochondrial iron transport via MFRN1 is required for erythroid cell cycle progression

**DOI:** 10.1101/2022.08.22.504833

**Authors:** Mark Perfetto, Aidan Danoff, Muhammad Ishfaq, Heidi Monroe, Aiden Mohideen, Meilin Chen, Amber N. Stratman, Satoshi Okawa, Yvette Y. Yien

## Abstract

Iron metabolism drives key erythropoietic processes, including hemoglobinization, survival, and proliferation. Here, we developed *in vivo* methods to interrogate how iron regulates erythropoiesis and report that mitochondrial iron transport via MFRN1 is essential for erythroid cell cycle progression. *mfrn1* embryos had severely decreased erythroid cell number caused by cell cycle arrest at G2/M. They had enlarged nuclei, suggesting a mitotic defect. Iron supplementation rescued the cell cycle defect, implicating mitochondrial iron deficiency as its cause. In contrast, *fpn1* mutants, anemic from systemic iron deficiency, had less severe decreases in erythroid mitochondrial iron than *mfrn1* mutants and no proliferative defects. scRNAseq and FACS analyses for *cd41* (thrombocytic) and *gata1* reporters indicated that developmental defects in *mfrn1* mutants were largely erythroid restricted. *mfrn1* mutant *gata1*^*+*^ erythroid progenitors were severely decreased at 3 dpf, and a further decrease in *globin-* expressing terminally differentiating erythroid cells. While wild-type erythroid cells mostly lost expression of the *gata1* progenitor marker by 3 dpf, *mfrn1* mutant erythroid cells retained *gata1* expression. These data are consistent with a model where mitochondrial iron transport facilitates development of *gata1*^*+*^ erythroid progenitors and is required for the completion of erythropoiesis by facilitating mitosis in the terminal cell cycles.

**Key points:**

- Mitochondrial iron import through MFRN1 is required for progress through the G2/M checkpoint in the terminal cell cycles leading to terminal erythroid differentiation.
- Mitochondrial iron supply through MFRN1 is required for maintenance/proliferation of erythroid progenitors and terminal maturation into globin expressing erythroid cells.

## Introduction

Iron deficiency anemia is a formidable public health problem, afflicting 27% of the world’s population and accounting for 8.8% of the world’s disability^1-3^. In the US, up to 77% of 12-21-year-old females are iron deficient^4-10^. Loss-of-function studies demonstrate erythropoietic defects including decreased cell number, morphological and enucleation defects caused by iron metabolism defects^11-16^. While iron metabolism pathways are key regulators of erythropoiesis, their mechanisms remain poorly understood.

Mitochondria are the site of eukaryotic heme and iron-sulfur (Fe-S) cluster synthesis. Mitochondrial iron processing and cytosolic export are required for iron incorporation into cytosolic apo-proteins ^17^. The mitoferrins are the main importers of mitochondrial iron in eukaryotes^18^. Loss of function mutations in vertebrate MFRN1 significantly decreased heme synthesis, Fe-S cluster formation and assembly of mitochondrial iron binding proteins^8,19-22^. Whole body *Mfrn1* mouse and zebrafish loss-of-function caused severe anemia resulting in embryonic lethality, while an adult hematopoietic-specific deletion of *Mfrn1* was anemic with increased numbers of erythroblasts and erythroid progenitors^7^.

This study dissects the role of mitochondrial iron during erythropoiesis and determines that mitochondrial iron deficiency causes erythropoietic defects distinct from systemic iron deficiency. We used zebrafish as an *in vivo* model for several reasons. Zebrafish develop externally, allowing us to utilize EdU labeling for cell cycle analyses in live embryos. To confirm that the developmental defects were caused by iron transport defects, we performed iron supplementation experiments. The external development of zebrafish embryos enabled us to treat them with lipophilic iron chelates ^18^ which is not feasible in mammalian fetuses which develop *in utero*. Due to the small number of cells in early zebrafish embryos, the field has typically relied on sorting embryos into “mutant” and “wildtype and heterozygote” groups for fluorescence-activated flow sorting (FACS) experiments based on visible phenotypes, which precludes interrogating subtler, potentially clinically relevant heterozygote phenotypes. We have therefore designed methods to label and sort erythroid cells from single zebrafish for imaging, mitochondrial iron content, and cell cycle analyses, which provide a resource to the zebrafish field, substantially increasing the resolution of data acquisition from live organisms.

Here, we interrogate erythropoietic defects in the absence of MFRN1 to model erythroid mitochondrial iron deficiency^8^, or FPN1, the transporter of iron from the gut or yolk syncytia into the organism, as a model for systemic iron deficiency^9,23,24^. We demonstrate that mitochondrial iron transport via MFRN1 is required for the development and terminal differentiation of erythroid progenitors. In its absence, erythroid progenitors are arrested at the G2/M cell cycle checkpoint, precluding completion of proliferative cycles required for maturation. In contrast, FPN1-deficient erythroid cells which have a systemic iron deficiency, did not experience these proliferative defects, likely due to higher mitochondrial iron levels. We demonstrate that mitochondrial iron transport plays a specific role during terminal erythropoiesis.

## Methods and Materials

### Iron-hinokitiol treatment

Embryos were harvested at 0 dpf, dechorionated, and incubated with 10uM FAC (Fisher I72500) and 1uM hinokitiol (TCI H0142) or 20uM FAC and 15uM hinokitiol in DMSO in E3 media^18^. Media was changed daily.

### FACS sorting of zebrafish erythroid cells onto slides

Zebrafish embryos were washed with 1% BSA in 100uL PBS, pH 7.4 (PBSB) and dissociated. 400uL PBSB was added to the suspension, strained with a 70um filter and centrifuged in a swinging bucket rotor for 5 minutes at 300g at RT. Cells were fixed with 0.25% glutaraldehyde (GTA) in PBSB for 5 minutes at RT, washed in PBSB 3 times and filtered. Cells were sorted on a FACSAria Fusion. GFP^+^ cells were sorted at 488nm. Flow rate was set at 3-4. To sort cells onto slides, drop delay was adjusted each time to achieve more than 98% deflection. Cells were collected at a density of 100 cells/spot and dried overnight in the dark at RT.

### Analysis of mitochondrial iron content

Dissociated single zebrafish cells were stained with 10uM rhodamine B-[(1,10-phenanthroline-5-yl)-aminocarbonyl]benzyl ester (RPA) (Bionet TS-7614) in PBSB for 30 minutes at RT in the dark. The cells were then washed 3 times with PBSB and strained with a 70uM filter. FACS analysis was carried out on a FACS Aria. 488nm and 532nm lasers were used for GFP and RPA analysis respectively.

### Cell cycle analysis

Single zebrafish embryos were washed twice with PBSB and incubated in ice-cold 1.5mM EdU (Thermo C10419) in 15% DMSO, or 15% DMSO vehicle control in the dark, for 1 hr. The embryos were washed 2x with PBSB and incubated at 28°C for 30 minutes followed by dissociation. Single cells were then washed twice with PBSB and fixed in 2% paraformaldehyde (PFA) in the dark for 10 minutes at RT. Cells were washed 3 times in PBSB and subsequently permeabilized in 500uL 0.1% Triton X-100 in PBSB for 15 minutes at RT in the dark. Cells were washed 3x in PBSB and subjected to a Click-It reaction to attach a 694nm azide-fluorophore to the incorporated EdU following the manufacturers’ guidelines (Thermo C10419). Cells were washed 3x in PBSB. GFP^+^ cells were identified by immunofluorecence. Washed cells were incubated with 1:1000 anti-GFP (Abcam Ab290) in the dark for 30 minutes at RT, washed 3X with PBSB and then incubated with 1:800 anti-mouse-488 (Abcam Ab150077) in the dark for 30 minutes at RT. The cells were then washed 3 times in PBSB and mixed with 1:200 DyeCycle-Violet (Invitrogen V35003) and filtered through a 70um strainer. The cells were incubated in the dark for 30 minutes at RT and analyzed by FACS with a FACS Aria. 450/50nm, 488nm, and 635nm lasers were used for DyeCycle-Violet, GFP, 694-EdU analysis respectively.

### Statistical analysis

Statistical analysis was performed using Prism to conduct Tukey-Kramer analysis. Statistical significance was set at α= 0.05.

### Preparation of zebrafish cells for scRNAseq

10, 3-dpf zebrafish embryos from WT and *mfrn* mutant fish were homogenized in 300 µL PBS containing 10% FBS and through a 70 µm cell strainer. The cells/debris that were caught in the filter were genotyped (Supplemental Methods). The single cell suspension was filtered again through a 40µm cell strainer.

### scRNA-seq data analysis

10x scRNA-seq data processing and analyses were performed with the Seurat v.4.3.0 R package^25^. Low quality cells were excluded by filtering out cells with fewer than 1000 detected genes and cells with mitochondrial RNA more than 15%. The integration of the WT and *mfrn* mutant data was conducted using *FindIntegrationAnchors* function with default parameters. The data were then CPM-normalized and clustered using the *RunPCA, FindNeighbors* and *FindClusters* functions using the first 30 principal components at 0.8 resolution. The 2D projection of the clustering and visualization were carried out by the *RunUMAP* and *DimPlot* functions.

## Results

Previous studies have obtained embryonic blood by tail amputation^10^, sorting zebrafish embryos based on visible phenotypes to obtain sufficient numbers of cells for FACS analysis ^26^ or analyzing erythroid cells from adult zebrafish which have larger cell numbers ^8,27^. However, these approaches have several limitations. Hematopoietic mutants have severely decreased numbers of erythroid cells; further cell losses during tail amputation precludes acquisition of quantitative data. Many hematopoietic mutations are embryonic lethal, precluding collection of cells from the adult. Grouping zebrafish embryos into “wild-type/heterozygote” or “homozygous mutant” groups according to their visible phenotypes for batch processing ^26,28^ precludes correlating quantitative information with specific genotypes, particularly since subtle, potentially clinically relevant heterozygote phenotypes are visually indistinguishable from wild-type embryos. Lastly, grouping by visible phenotype masks statistical variability within individual genotypes, necessitating the development of methods to genotype and sort erythroid cells from single zebrafish embryos (Figure 1A)

**Figure 1.**
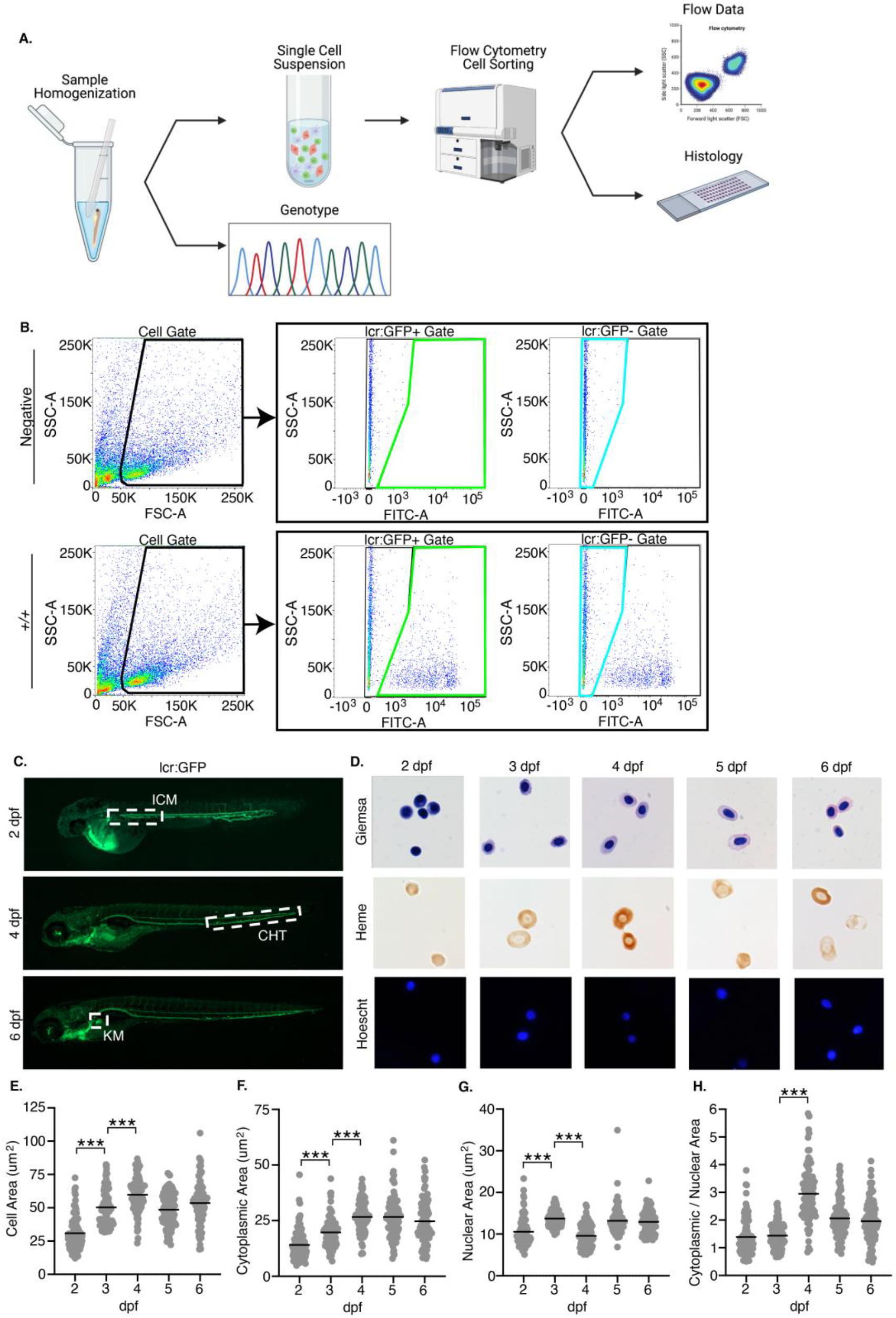
Experimental workflow and gating scheme used for sorting lcr:GFP^+^ erythroid cells. (A) Individual zebrafish are homogenized into a single cell suspension in PBS containing 1% BSA. Single cells were collected by filtration through a 70 µm cell strainer. Cells/debris caught in the filter were rinsed and collected for genotyping analysis by PCR and Sanger sequencing of the PCR product. The filtrate (single cell suspension) was processed for FACS analysis and sorting. When indicated, cells were sorted onto microscope slides and processed for histological analysis. (B) Gating scheme to sort lcr:GFP^+^ and lcr:GFP^-^ cells. Gates were set up using wild-type AB zebrafish embryos as a negative control. The single cell population is gated by FSC and SSC (left panels; black outline). The GFP^-^ population is defined by the FITC signal on cells from the negative controls which do not express GFP (teal outline); any FITC signal above that is defined as GFP^+^ (green outline). (C) lcr:GFP marks erythroid cells during development. At 2 dpf, primitive erythroid cells emerge from the intermediate cell mass (ICM); at 4 dpf, definitive erythroid cells emerge from the caudal hematopoietic tissue (CHT). By 6 dpf, the kidney marrow (KM) becomes the primary site of erythropoiesis. From 2 dpf onwards, circulating erythroid cells (green) are also found in blood vessels. (D) Giemsa, Benzidine and Hoechst staining of sorted lcr:GFP^+^ erythroid cells from pooled zebrafish embryos. Erythroid cells are hemoglobinized by 2 dpf, but acquire a more elliptical shape at about 4 dpf. (E) Erythroid cells grow in area through 4 dpf, largely due to increase in (F) cytoplasmic area. (G) Nuclear area is mostly constant from 2-6 dpf. (H) Cytoplasmic/nuclear area markedly increased about 4 dpf, attributable to the increase in cytoplasmic area around this time. Magnification: 63x. *** indicates p<0.001

To characterize erythroid cells from single zebrafish embryos, we used the Tg(*lcr:eGFP*) transgenic line which expresses GFP under the control of the globin locus control region, marking erythroid cells GFP with expression ^29^ (gating strategy in Figure 1B). From 2 dpf, we observed GFP^+^ erythroid cells in circulation (Figure 1C). To validate this approach, we fixed and sorted GFP^+^ cells from multiple 2-6 dpf zebrafish embryos onto microscope slides, subjecting cells to Benzidine/Hoechst or Wright-Giemsa staining for morphological and heme analysis. ImageJ analysis indicated that erythroid cell size grew and become more elliptical from 2-4 dpf (Figure 1E), predominatly driven by increases in cytoplasmic area (Figure 1F) ^30-32^. The sizes of erythroid nuclei were largely fixed by 2 dpf and did not condense from 2-6 dpf, unlike mammalian erythroid cells whose nuclei condense during terminal differentiation (Figure 1G). This suggests that condensation of zebrafish erythroid nuclei and nuclear size determination occurs during early differentiation, unlike in vertebrates ^33-35^. The morphological changes are reflected in increased cytosolic/nuclear area between 2-4 dpf (Figure 1H). These data show that our method of FACS sorting and immobilizing erythroid cells onto microscope slides recapitulates observations from published zebrafish erythroid cell isolation methods.

To interrogate the role of mitochondrial iron transport in erythropoiesis, we compared the erythropoietic phenotypes of *frascati* (*frs;* Mfrn1) and *weissherbst* (*weh;* Fpn1) zebrafish mutants previously identified in screens^8,23^. MFRN1 is the key mitochondrial iron transporter for erythroid heme and iron-sulfur cluster synthesis ^7,8,19^. FPN1 transports iron from the yolk to the embryo during early development^23,24^, hence *weh* embryos generally model systemic iron deficiency. We crossed *frs* (Mfrn1) and *weh* (Fpn1) heterozygotes with *Tg (globin lcr:eGFP)* transgenic zebrafish to dissect their erythropoietic phenotypes. To characterize mitochondrial iron content, we stained 3 dpf wild-type and *frs* or *weh* mutant; *Tg (lcr:eGFP)* zebrafish embryos with rhodamine B-[(1,10-phenanthrolin-5-yl)aminocarbonyl]benzyl ester (RPA), an indicator that localizes to the mitochondria whose fluorescence is quenched by labile Fe^2+^ (the redox state in heme and Fe-S synthesis)^36^. RPA fluorescence in cells from single embryos was quantified by FACS. Consistent with the role of *Mfrn1* in erythroid mitochondrial iron import, *frs/frs* erythroid cells exhibited over 10-fold increased RPA fluorescence, indicating drastically decreased mitochondrial Fe^2+^ (Figure 2A).

**Figure 2.**
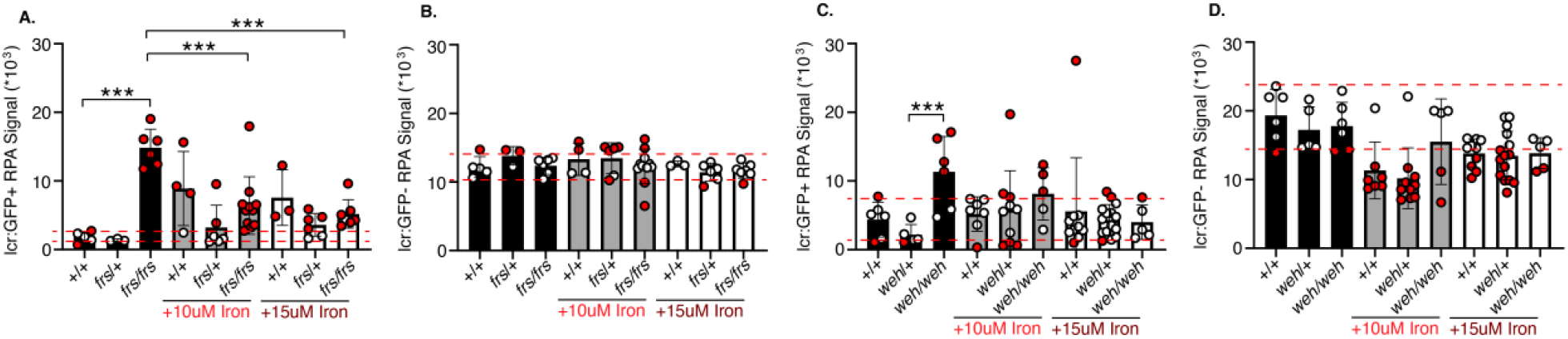
RPA staining of live zebrafish embryos confirms decrease of mitochondrial iron in erythroid cells of mutant zebrafish lines and ability of iron-hinokitiol to increase erythroid mitochondrial Fe^2+^ levels in mutant fish. (A) *frs/frs* mutant lcr:GFP^+^ erythroid cells have an increase in RPA signal, indicating significantly decreased Fe^2+^ levels relative to wild-type cells. Addition of iron-hinokitiol decreased RPA signal, indicating increased mitochondrial Fe^2+^. (B) *frs/frs* mutant lcr:GFP^-^ (non-erythroid) cells did not significantly differ from wild-type cells in their RPA signal, indicating that *Mfrn1* was largely not required for maintenance of non-erythroid mitochondrial Fe^2+^ levels. Addition of supplemental iron did not alter Fe^2+^ levels in *frs/frs* non-erythroid cells. (C) *weh/weh* mutant lcr:GFP^+^ erythroid cells have increased RPA signal, indicating significantly decreased Fe^2+^ levels relative to *weh/+* cells. Addition of iron-hinokitiol decreased RPA signal, indicating increased mitochondrial Fe^2+^. (D) *weh/weh* mutant lcr:GFP^-^ (non-erythroid) cells did not significantly differ from wild-type cells in their RPA signal, indicating that *Fpn1* (despite its requirement for embryonic utilization of yolk iron) was largely not required for maintenance of non-erythroid mitochondrial Fe^2+^ levels. Addition of supplemental iron did not alter Fe^2+^ levels in *weh/weh* non-erythroid cells. *** indicates statistical significance by the Tukey Kramer test at 95% significance. Dashed red lines indicate 90% confidence intervals as defined by wild-type controls; red data points indicate data points that fall outside these 90% confidence intervals.

Iron supplementation is a first-line treatment for iron deficiency, and Fe^3+^ chelates (since transferrin binds Fe^3+^) are used to model pharmacologic iron supplementation ^5,18,37^. Hence, we asked if Fe (ferric ammonium citrate)-hinokitiol, a lipophilic iron chelate that transports iron across membranes, could restore mitochondrial Fe^2+^ levels *in vivo*. This is a critical control because lipophilic Fe^3+^ bypasses transferrin-mediated iron uptake which reduces Fe^3+^ to Fe^2+^ in the acidified endosome via STEAP proteins ^38^. We treated zebrafish embryos with 10 or 15 µM Fe-hinokitiol (Supplemental Figure 1) which caused significantly decreased RPA fluorescence, indicating effective reduction of Fe^3+^-hinokitiol to Fe^2+^ (Figure 2A). Non-erythroid wild-type, *frs/+* and *frs/frs* cells had similar mitochondrial Fe^2+^ levels after Fe-hinoktiol treatment, indicating that responses to iron supplementation are tissue-specific (Figure 2B).

We observed that the mitochondrial iron defect in *weh/weh* mutant erythroid cells was less severe than that of *frs* mutants, with their RPA staining about double that of wild-type controls (Figure 2C). This is likely because of the presence of MFRN1, or because erythroid FPN1 deficiency increases iron retention^39^. Similar to *frs/frs* non-erythroid cells, mitochondrial Fe^2+^ content in *weh/weh* non-erythroid cells did not significantly differ from wild-type controls (Figure 2D). Although *weh/weh* embryos model systemic iron deficiency, these data reflect that erythroid iron deficiency is more pronounced because of increased iron demand ^39^. In Figures 2A and C, we also noted that iron doses that were effective for restoration of erythroid mitochondrial iron also caused a paradoxical *decrease* in WT mitochondrial iron levels, possibly indicating that these doses cause toxicity (Supplemental Figure 2).

We sorted and imaged erythroid cells from 3 dpf single embryos from *frs/+; Tg (globin lcr:eGFP)* incrosses (gating strategy in Figure 1). Here, we analyzed potentially clinically significant phenotypes by delineating 90% confidence intervals on the wild-type population as this is the method used to define reference ranges in clinical laboratory tests^40,41^, in addition to analyses of statistical significance. We obtained high quality sequencing data that distinguished between genotypes (Figure 3A). Despite small numbers of *frs/frs* erythroid cells (Figure 3B), we were able to sort them onto slides for morphological and heme staining. Consistent with previous work, they were poorly hemoglobinized and had much larger nuclei than wild-type and heterozygote cells (Figure 3A). To determine if these phenotypes could be rescued by iron supplementation, we treated embryos with 10 or 15 µM Fe-hinokitiol^18^. Treatment with 15 µM iron-hinokitiol increased *weh/weh* erythroid mitochondrial Fe^2+^ to wild-type levels. Iron supplementation increased hemoglobinization and decreased the cell and nuclear size of *frs/frs* erythroid cells (Figure 3A, Supplemental Figures 2-4). The proportion of GFP^+^ erythroid cells in *frs/frs* embryos was significantly decreased compared to wild-type and heterozygote siblings; this was increased by iron (Figure 3B). This decrease in erythroid cell number occurred relatively later in erythroid development; at 1.5 dpf, we did not yet observe a decrease in the *lcr:* GFP^+^ erythroid population (Supplemental Figure 5). FACS analysis showed *frs/frs* erythroid cells were significantly larger than wild-type or *frs/+* cells, and cell size was decreased by iron supplementation (Figure 3C). Benzidine staining indicated significantly decreased *frs/+* erythroid hemoglobinization, and was more pronounced in *frs/frs* cells (Figure 3D). The decreased hemoglobinzation in *frs/+* heterozygotes was not visibly detectable when screening whole embryos (Supplemental Figure 2) but many erythroid cells had benzidine staining intensity below the lower bound of the 90% confidence intervals set by wild-type erythroid cells ^40,41^. These data highlight the importance of quantitating subtle phenotypes in heterozygote animals as these may be clinically relevant. Iron supplementation significantly increased *frs/frs* erythroid hemoglobinization but not *frs/+* erythroid cells, perhaps indicating homeostatic mechanisms that prevent cells from acquiring excessive iron (Figure 3D). *frs/frs* erythroid cells were much larger than wild-type controls, and many *frs/+* nuclei were larger than the 90% confidence intervals of wild-type nuclei (Figure 3D), significantly decreasing cytoplasmic/nuclear ratio (Figure 3E). Iron supplementation significantly decreased the sizes of *frs/frs* erythroid nuclei (Figure 3D), normalizing their nuclear/cytoplasmic ratio (Figure 3E).

**Figure 3.**
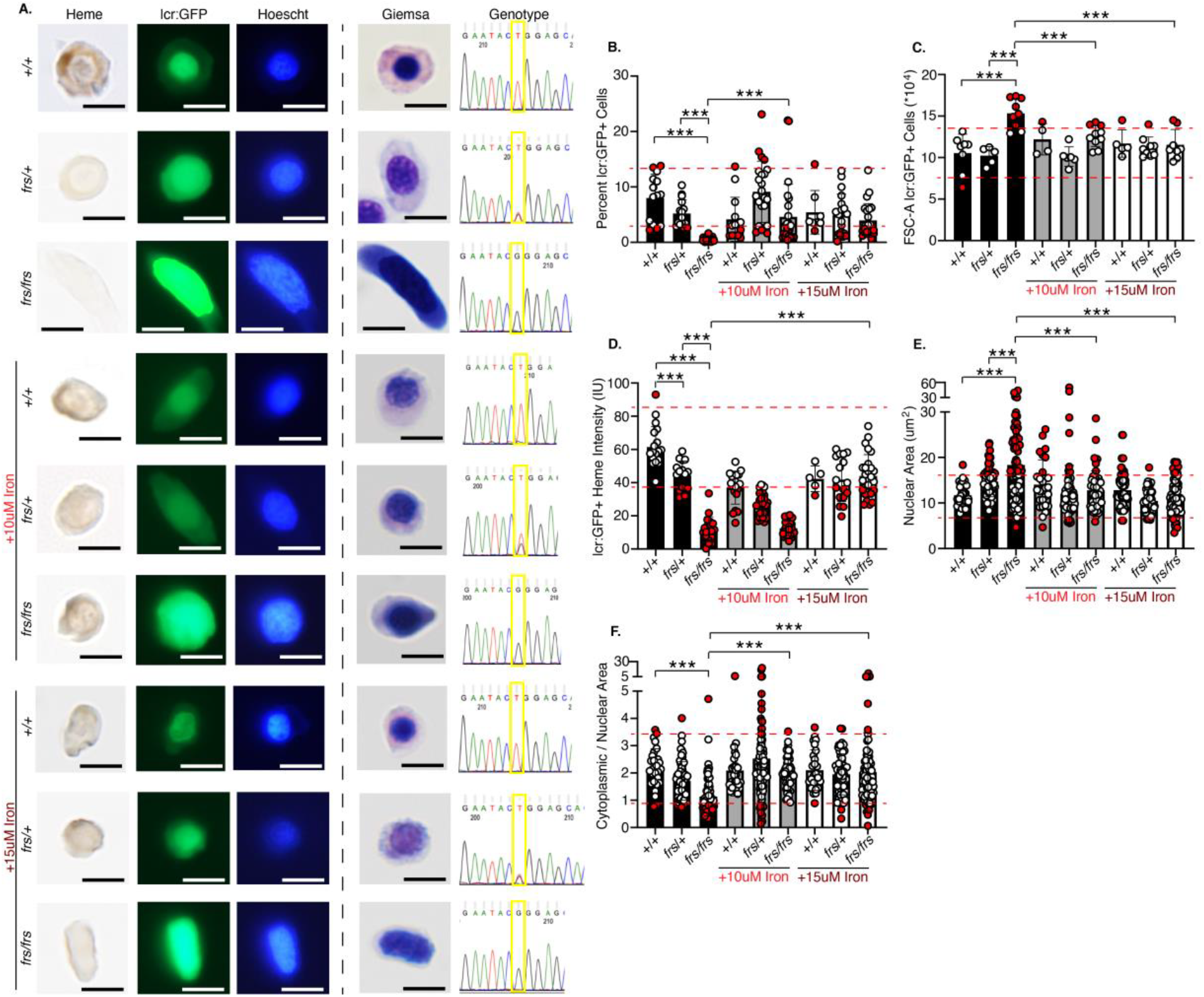
Sorting, imaging and genotyping of single 3 dpf zebrafish WT, *frs/+* and *frs/frs* embryos demonstrates feasibility of sorting erythroid cells from single embryos; iron supplementation significantly ameliorates the effects of *mfrn1* deficiency. (A) Sorting of GFP^+^ erythroid cells from single zebrafish embryos onto slides and followed by benzidine and Hoechst, or Giemsa staining. Genomic DNA was extracted from unsorted cells/debris and subjected to PCR with primers flanking the mutation region; the isolated PCR product was sequenced. Heterozygotes had overlapping nucleotide peaks at the mutation site. GFP^+^ WT cells were benzidine stained, indicating that sorted cells were from the erythroid lineage; iron treatment showed some evidence of toxicity (wrinkled morphology, decreased benzidine staining of individual cells). Both Hoechst and Giemsa staining revealed that *frs/frs* erythroid cells had very enlarged nuclei. This phenotype was largely reversed by iron supplementation. (B) Both *frs/+* and *frs/frs* mutant embryos had a significantly decreased number of erythroid cells. Erythroid cell proportion in *frs/frs* were significantly increased by iron treatment. (C) FSC analysis indicates that *frs/frs* erythroid cells are significantly larger than WT and *frs/+*. This abnormality was largely reversed by iron supplementation. (D) ImageJ analysis of benzidine stained cells indicated a significant decrease in heme staining in both *frs/+* and *frs/frs* erythroid cells (see Supplemental Figure 3). 15 uM iron supplementation was required for restoration of hemoglobinzation levels to WT. (E) ImageJ analysis of Giemsa stained cells revealed that *frs/+* and *frs/frs* erythroid cells had enlarged nuclei. These phenotypes could be reversed by iron supplementation. (F) The ratio of cytoplasmic/nuclear area was decreased in *frs/+* and *frs/frs* erythroid cells and these phenotypes could be reversed by iron supplementation though cell shapes qualitatively remained more variable than wild-type cells (see Supplemental Figure 4). *** indicates >95% significance.

Collectively, these data indicate that the *frs/frs* erythropoietic defects result from defective mitochondrial iron transport. Is this erythropoietic defect specific to MFRN1 loss of function? To address this, we investigated erythroid differentiation in *weh* (Fpn1) mutant; *Tg (globin lcr:eGFP)* zebrafish which mimic systemic iron deficiency ^9,23^. *weh/weh* erythroid cells had severe hemoglobinization defects. However, their cell and nuclear sizes were not larger than wild-type and heterozygote cells (Figure 4A; Supplemental Figures 6 and 7). Unlike *frs/frs* embryos, *weh/weh* embryos did not have a significant decrease in erythroid cell numbers (Figure 4B). *weh/weh* erythroid cells were similar in size to wild-type and heterozygote cells (Figure 4C). We supplemented wild-type and *weh/weh* zebrafish with Fe-hinokitiol^18^; 15 µM of iron-hinokitiol restored hemoglobinization to wild-type levels (Figure 4D; representative whole mount embryo Supplemental Figure 2B). The nuclear areas of *weh/weh* erythroid cells were significantly smaller than wild-type cells which was reversed by 15µM Fe-hinokitiol (Figure 4E) though cytoplasmic:nuclear ratios were similar across genotypes (Figure 4F).

**Figure 4.**
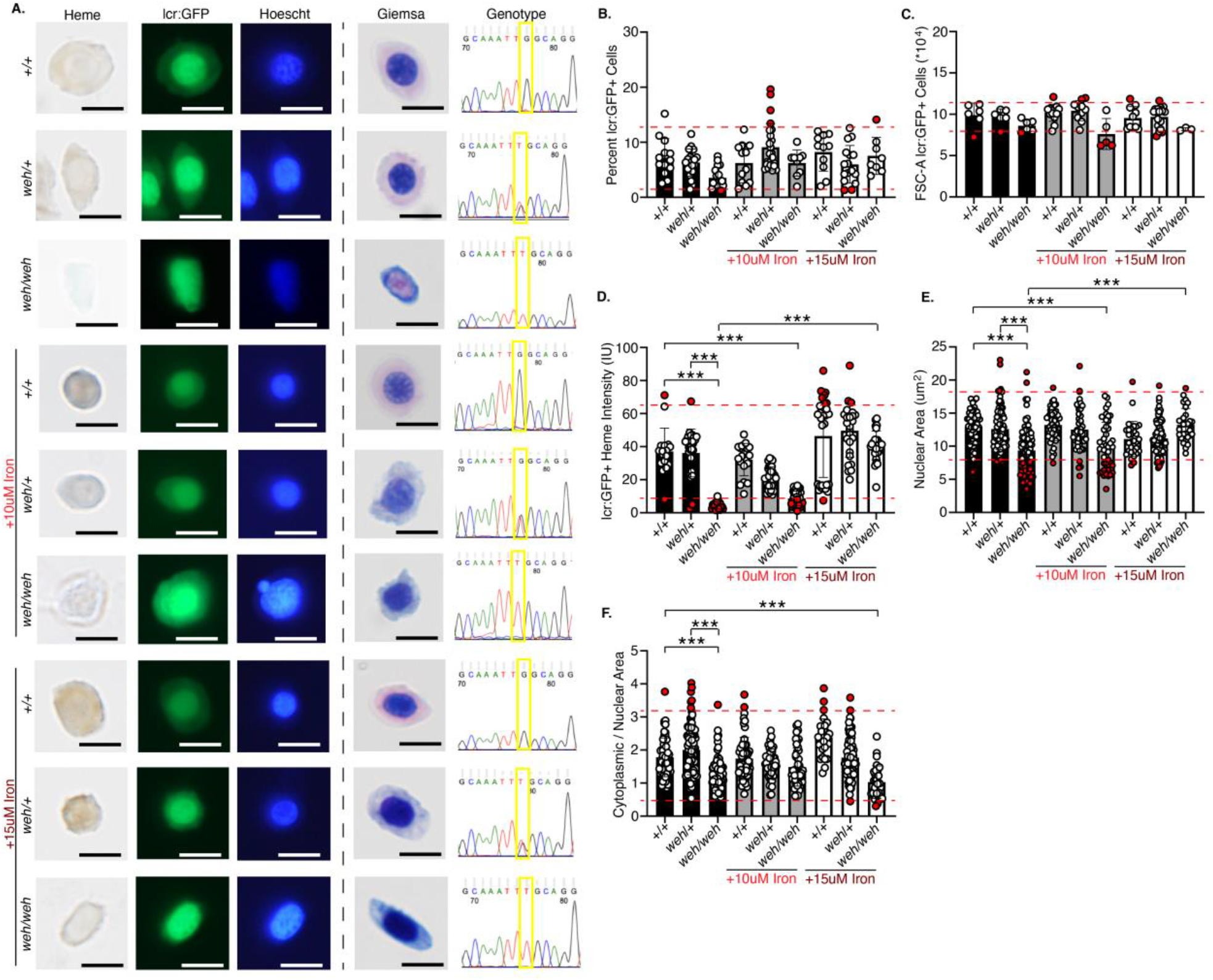
Sorting, imaging and genotyping of single 3 dpf zebrafish WT, *weh/+* and *weh/weh* embryos demonstrates feasibility of sorting erythroid cells from single embryos; iron supplementation ameliorates the effects of *Fpn1* deficiency. (A) Sorting of GFP^+^ erythroid cells from single zebrafish embryos onto slides and followed by benzidine and Hoechst, or Giemsa staining. Both Hoechst and Giemsa staining revealed that *weh/weh* erythroid cells were heme deficient and smaller than WT cells. This phenotype was largely reversed by iron supplementation. (B) *weh/weh* mutant embryos did not have significantly decreased numbers of erythroid cells at 72 hpf. (C) FSC analysis confirmed that *weh/weh* erythroid cells are significantly smaller than WT and *weh/+*. This abnormality was reversed by iron supplementation. (D) ImageJ analysis of benzidine stained cells indicated a significant decrease in heme staining in *weh/weh* erythroid cells. 15 uM iron supplementation was required for restoration of hemoglobinzation levels to WT levels (Supplemental Figure 6) (E) ImageJ analysis of Giemsa stained cells revealed that *weh/weh* erythroid cells had smaller nuclei than WT, which was corrected by iron supplementation. (F) The ratio of cytoplasmic/nuclear area was decreased in *weh/weh* erythroid cells and these phenotypes could not be reversed by iron supplementation, and cell morphology remained qualitatively different from WT cells (see Supplemental Figure 7). Scale bar = 5um. *** indicates >95% significance by the Tukey Kramer test.

While both *frs* (*Mfrn1*) and *weh* (*Fpn1*) mutants exhibited hemoglobinization defects, the *frs/frs* mutants exhibited more severe decreases in erythroid cell number than the *weh/weh* mutants. Because erythroid development is intricately co-regulated with cell cycle progression ^42-46^, we examined the roles for *Mfrn1* and *Fpn1* in cell cycle regulation. We incubated whole 3 dpf embryos in 5-ethynyl-2’deoxyuridine (EdU), a thymidine analog incorporated into DNA during the cell cycle S phase and subsequently attached an azide fluorophore in EdU labeled cells. Cells were stained with DyeCycle Violet stain for DNA content measurements. Since attachment of the fluorophore to EdU quenched GFP fluorescence, we labeled GFP^+^ cells with an anti-GFP primary antibody and an Alexa 488 conjugated secondary antibody. The DyeCycle^lo^ EdU^neg^ population was gated as G0/G1, while the DyeCycle^hi^Edu^neg^ population was gated as G2/M. The EdU^pos^ population was defined as S phase (Supplemental Figure 8).

Earlier in development at 1.5 dpf, when there was not yet an erythroid cell number defect in *frs/frs* embryos (Supplemental Figure 5), we did not observe cell cycle defects in the GFP^+^ erythroid population (Supplemental Figure 9). We then carried out similar cell cycle analyses on 3 dpf embryos in which *frs* mutant embryos had observable erythropoietic defects. At 3 dpf, the number of GFP^+^ (erythroid) cells with sub-G0 DNA did not significantly differ between genotypes (Figure 5A). Most 3 dpf wild-type erythroid cells were in G0/G1, in contrast to erythroid cells earlier in development at 1.5 dpf, which were more proliferative (Supplemental Figures 9, 11). While there were not statistically significant differences between the proportion of WT and *frs/+* erythroid cells in G0/G1, most *frs/+* embryos had proportions of G0/G1 erythroid cells below the 90% confidence intervals defined by wild-type embryos. There was a significant decrease in erythroid cells in G0/G1 in *frs/frs* embryos, suggesting that they had exited quiescence. This was reversed by iron supplementation (Figure 5B). We did not observe changes in the proportions of *frs/frs* mutant erythroid cells in S phase (Figure 5C), but the proportion of *frs/+* erythroid cells in G2/M had doubled and most *frs/frs* erythroid cells were in G2/M suggesting cell cycle arrest, corresponding with decreased erythroid cell number (Figure 3). Iron supplementation significantly decreased the proportion of *frs/frs* erythroid cells in G2/M, suggesting release from cell cycle arrest (Figure 5D), corresponding to increased erythroid cell numbers upon iron treatment (Figure 3B).

Proportions of sub-G0 non-erythroid cells did not differ between genotypes (Figure 5E). *frs/frs* embryos had decreased numbers of non-erythroid G0/G1 cells that were increased by iron treatment, suggesting that some non-erythroid cell types require MFRN1 for iron transport and cell cycle regulation (Figure 5F). They did not experience S phase defects (Figure 5G), but had increased proportions of cells in G2/M, reversed by 10 µM Fe-hinokitiol. 15 µM increased percentages of G2/M WT and *frs/+* cells, suggesting toxicity of excess iron (Figure 5H).

**Figure 5.**
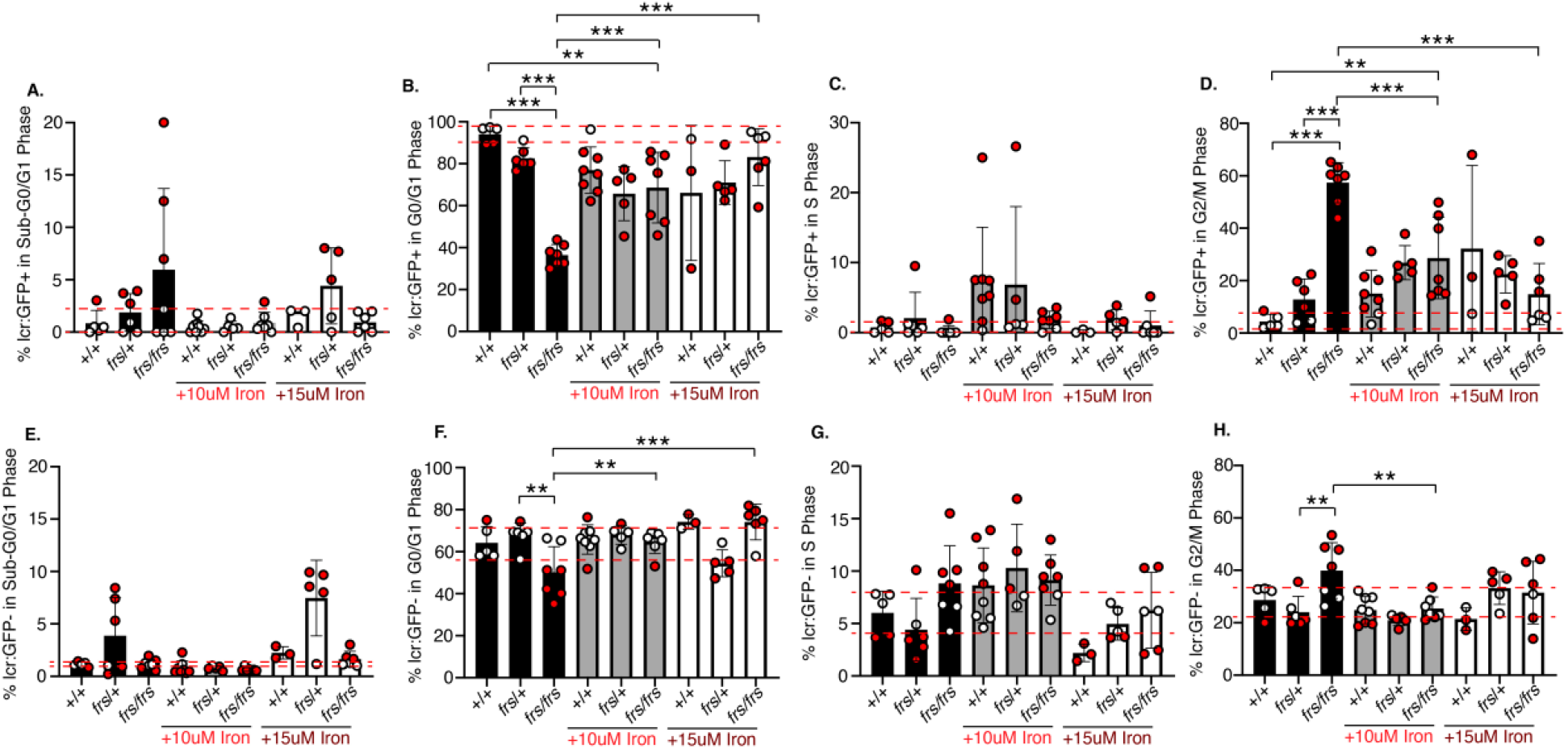
Iron transport via Mfrn1 is required for erythroid cell cycle progression. (A) Mfrn1 deficiency does not cause increased erythroid cell death. We did not observe a significant difference between WT, *frs/+* and *frs/frs* groups in the numbers of cells with sub-G0 DNA. (B) Most WT erythroid cells were in G0/G1, suggesting they were quiescent. There was a significant decrease in the numbers of *frs/+* and *frs/frs* erythroid cells in G0/G1. Addition of supplemental iron significantly increased the percentage of of *frs/frs* erythroid cells in G0/G1. (C) There were no significant differences between groups in the percentage of erythroid cells in S phase. (D) *frs/+* and *frs/frs* mutants had a significant increase in the percentage of erythroid cells arrested in G2/M. Addition of supplemental iron significantly reduced the percentage of mutant erythroid cells in G2/M arrest. However, even iron supplementation could not reduce the percentage of erythroid cells in G2/M to wild-type levels. (E) Mfrn1 deficiency did not significantly affect cell death in non-erythroid (GFP^-^) cells. (F) Mfrn1 deficiency significantly decreased the proportion of non-erythroid cells in G0/G1, suggesting an increase in the number of cells exiting quiescence. Supplemental iron restored the proportion of cells in G0/G1 to wild-type levels. (G) There were no significant differences between groups in the percentage of erythroid cells in S phase. (H) *frs/frs* mutants had a significant increase in the percentage of non-erythroid cells arrested in G2/M. Addition of supplemental iron significantly reduced the percentage of *frs/frs* non-erythroid cells in G2/M arrest to wild-type levels. ** indicates p<0.01, *** indicates p<0.001 by Student’s t-test; ** and *** obtained for pairs with 95% significance by Tukey-Kramer test.

Does erythroid cell cycle arrest occur in systemic iron deficiency? We analyzed cell cycle status in GFP^+^ cells obtained from incrosses of *weh/+*; Tg(*lcr:GFP*) zebrafish. The average percentages of erythroid cells in sub-G0 did not differ between genotypes (Figure 6A). In contrast with the *frs* mutants, *weh* mutants did not have decreased G0/G1 GFP^+^ erythroid cells (Figure 6B). Similarly, percentages of erythroid cells in S phase (Figure 6C) and in G2/M (Figure 6D) were similar across genotypes. We also did not observe genotype-dependent differences in percentages of non-erythroid cells with sub-G0 DNA (Figure 6E), in G0/G1 (Figure 6F), S phase (Figure 6I) or G2/M (Figure 6H). Collectively, these data indicate that the erythropoietic defect in *frs/frs* mutants is caused by G2/M arrest and ameliorated by supplemental iron, indicating that mitochondrial iron transport via MFRN1 is required for cell cycle progression in differentiating erythroid cells.

**Figure 6.**
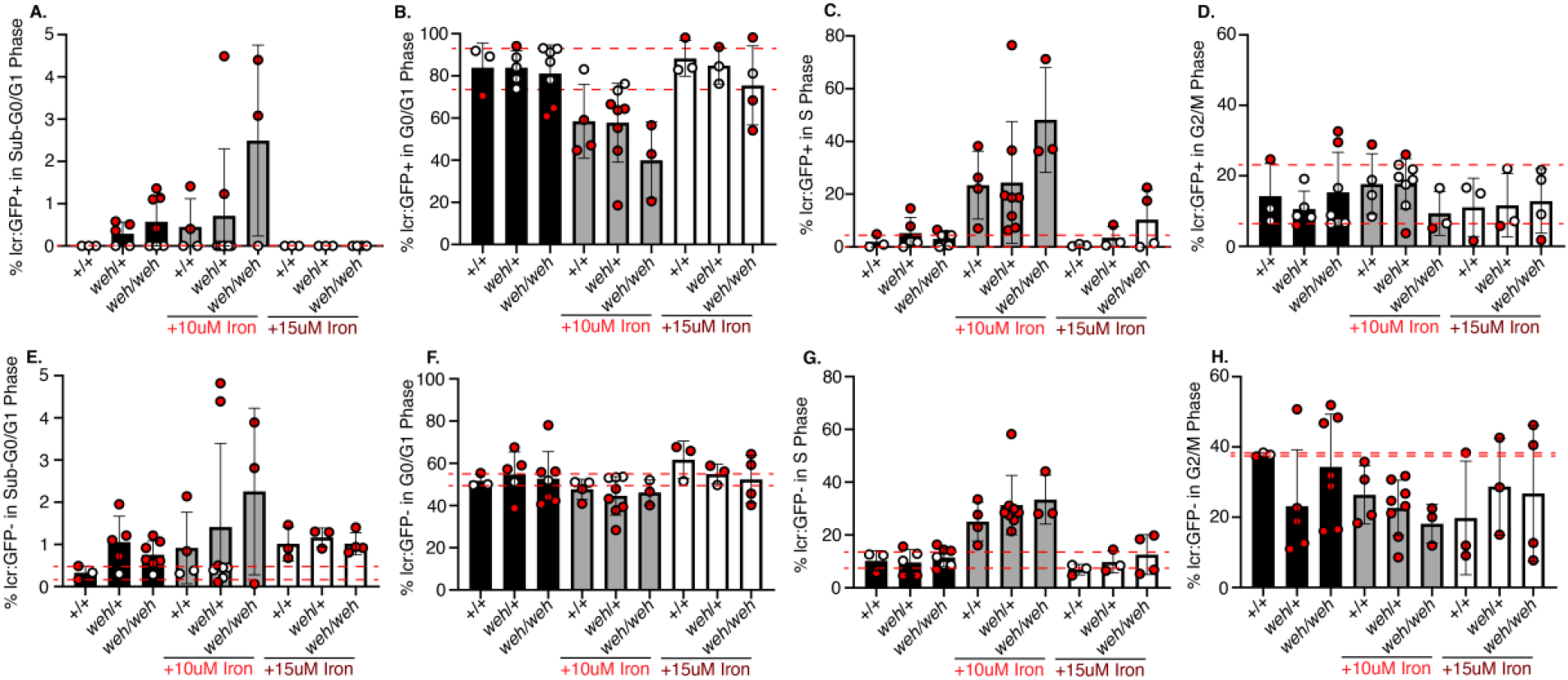
Fpn1 is not required for erythroid cell cycle progression even though homozygous mutants are deficient in mitochondrial iron (Figure 2). (A) Fpn1 deficiency does not significantly increase erythroid cell death (% cells with sub G0/G1 DNA), but there are increased numbers of *weh/+* and *weh/weh* embryos with increased numbers of % sub-G0 DNA relative to vehicle treated WT. Addition of 15 uM iron reduced cell death to WT levels. (B) *weh/+* and *weh/weh* mutants did not differ from the WT group in the number of erythroid cells in G0/G1. This remained consistent with iron supplementation (C) There were no significant differences between groups in the percentage of erythroid cells in S phase. (D) There were no significant differences between groups in the percentage of erythroid cells in G2/M. (E) Fpn1 deficiency increased cell death in non-erythroid (GFP^-^) cells. Addition of iron increased cell death in WT cells, so there were no differences in the % of non-erythroid cell with sub G0/G1 between WT and mutant groups. (F) Fpn1 deficiency did not significantly decrease the proportion of non-erythroid cells in G0/G1. (G) There were no significant differences between groups in the percentage of erythroid cells in S phase. (H) There were no significant differences between groups in the percentage of non-erythroid cells in G2/M.

Iron deficiency can decrease hematopoietic stem cell (HSC) ^47^ or megakaryocytic-erythroid progenitor (MEP) ^48^ commitment to the erythropoietic lineage by preferential expansion of HSCs^47^, or increased commitment of MEPs towards the megakaryopoiesis ^48^. We asked whether the erythropoietic defect in *Mfrn1* deficient animals^7,8^ was caused by altered hematopoietic lineage commitment. To address this, we conducted scRNAseq analysis of pooled WT and *frs/frs* embryos (genotypes in Supplemental Figure 10). Single cells obtained from 10 3 dpf embryos from each group were subjected to a 10x Genomics pipeline for scRNAseq analysis. The scRNA-seq data were processed and clustered using Seurat v.4.3.0 [29608179] at a resolution of 0.8 (Figure 7A) and clusters were annotated by expression of marker genes (Figure 7B; Supplemental Figure 11). Lineage defects in *Mfrn1* deficient embryos were predominantly restricted to the erythroid lineage (cluster 6), where *frs/frs* embryos had a large drop in the number of cells in this cluster with only 76 erythroid cells sequenced out 10,000 cells loaded (Figure 7A, Supplemental Table 1). GFP and globin expression was restricted to Cluster 6 (Figure 7B, Supplemental Table 2). Within cluster 6, approximately 1/3 of wild-type erythroid cells expressed *gata1a*, an early marker and master regulator of erythroid differentiation^49-51^, while over half of the *frs/frs* erythroid cells expressed *gata1a* (Figure 7B).

**Figure 7.**
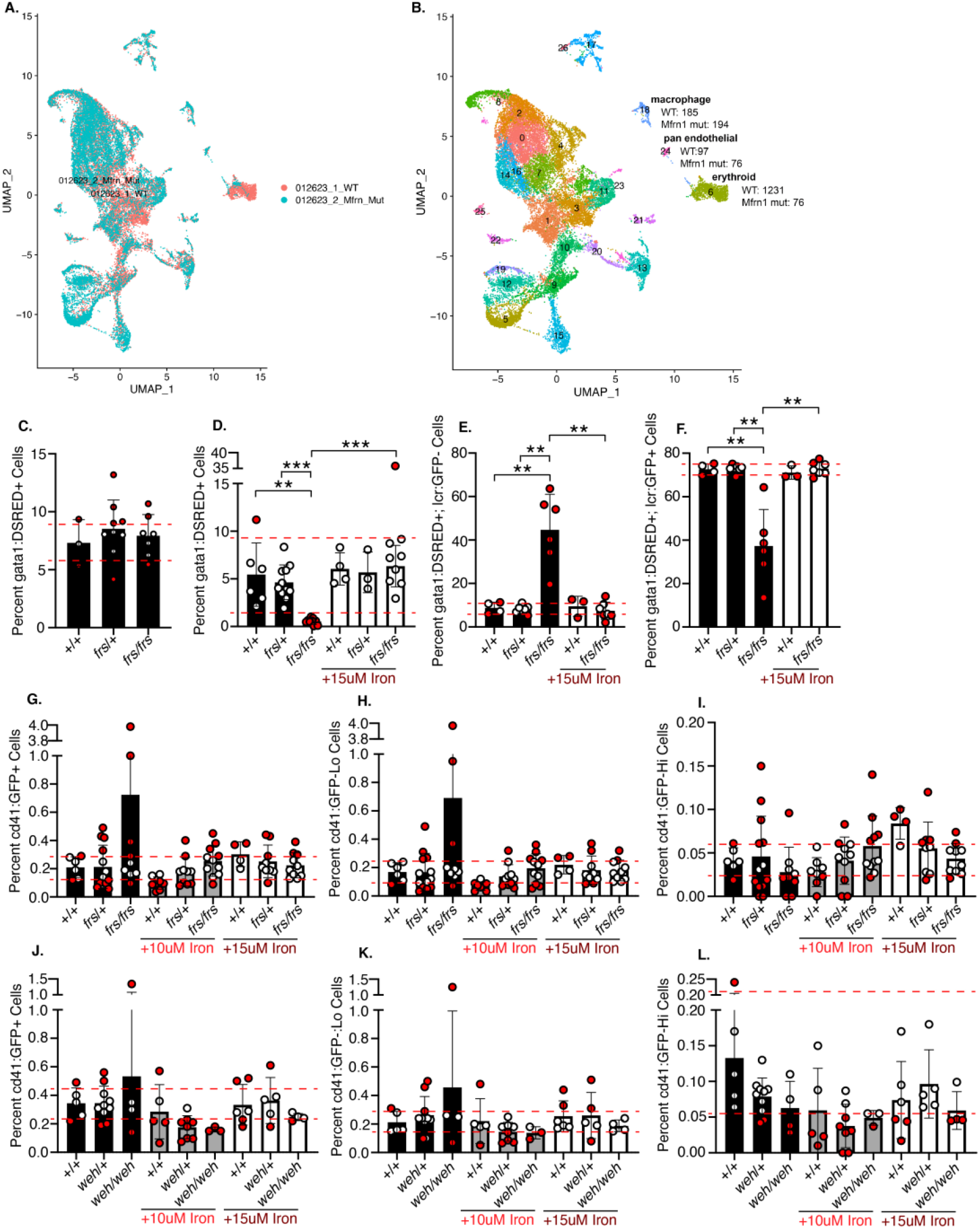
Mfrn1 is primarily required for erythroid cell maturation; neither *mfrn1 (frs)* nor *fpn1 (weh)* mutants have significant increases in the thrombocytic lineage that indicate lineage shifts in the MEP population. (A, B) Single cell RNAseq analysis indicates that the primary phenotype of *Mfrn1* deficiency is in formation of the mature erythroid lineage, which is *gata1a*^*-*^*/globin*^*+*^. There is also a significant defect in the development of gata1a^+^ erythroid cells in *frs/frs* mutants. (C) *frs/frs* mutants had normal proportions of *gata1a*^*+*^ cells at 24 hpf, indicating normal formation of erythroid precursors. (D) Vehicle-treated *frs/frs* mutants had a significant decrease in the proportion of *gata1a*^*+*^ cells at 72 hpf that was rescued to WT levels with 15uM iron-hinokitiol. (E, F) *frs/frs* mutant embryos have a significantly higher proportion of *gata1a*^*+*^;*lcr*^*--*^cells and lower proportion of *gata1a*^*+*^;*lcr*^*+*^ cells at 72hpf that is rescued to WT levels with 15uM iron-hinokitiol. (G) At 96 hpf, there was no significant difference in the proportion of CD41^+^ cells between vehicle-treated WT, *frs/+*, or *frs/frs* embryos. (H) *frs* and *frs/frs* mutants did not experience alterations in the development of cd41^lo^ cells, indicating that development of the adult hematopoietic stem cell population was unaltered. (I) *frs* and *frs/frs* mutants did not experience alterations in the development of cd41^hi^ cells, indicating that development of the thrombocyte population was unaltered. (J) At 96 hpf, *fpn1* mutation status or iron treatment did not alter percentages of total cd41^+^ cells. (K) *Fpn1* mutants did not experience alterations in the development of cd41^lo^ cells, indicating that development of the adult hematopoietic stem cell population was unaltered. (L) *Fpn1* mutants did not experience alterations in the development of cd41^hi^ cells, indicating that development of the thrombocyte population was unaltered. ** indicates p<0.01, *** indicates p<0.001 by Student’s t-test; ** and *** obtained for pairs with 95% significance by Tukey-Kramer test.

While GATA1 is required for erythroid specification and initiation of erythropoiesis, GATA1 downregulation is essential for completion of terminal erythropoiesis ^50,52^. The retention of *gata1a* expression in *frs/frs* erythroid cells is therefore indicative of arrested erythropoiesis.

To determine if erythropoietic commitment was defective in *frs/frs* mutants, we crossed *frs* mutants with *Tg*(*gata1:dsRed*) ^53^ zebrafish and quantitated dsRed fluorescent cells at 1 dpf, when *gata1* is largely expressed by erythroid progenitors ^54^. We found no significant differences in the numbers of dsRed^+^ cells between genotypes, suggesting that *Mfrn1* is not required for erythroid specification (Figure 7C), consistent with *Mfrn1* being a downstream target of GATA1 transcription^55^.

To determine if *Mfrn1* deficiency favored proliferation of *gata1*^*+*^ progenitors at the expense of terminal erythropoiesis, we crossed *frs* mutants with *Tg*(*gata1:dsRed*) and *Tg*(*lcr:GFP*) fish. At 3 dpf, *frs/frs* embryos had significantly decreased percentages of *gata1* - expressing cells compared to WT and *frs/+* embryos (Figure 7D). These data suggested that *frs/frs gata1a*^+^ erythroid progenitors did not undergo compensatory expansion at the expense of terminal erythropoiesis. The proportion of *gata1-*expressing cells was rescued by addition of 15 µM Fe-hinokitiol (Figure 7D). Most 3 dpf WT and *frs/+ gata1-*expressing cells were *lcr:GFP*^*+*^, indicating that *gata1* is primarily expressed in committed erythroid cells. In contrast, more than 40% of *frs/frs gata1*^*+*^ cells were *lcr:GFP*^*-*^, indicating developmental arrest at a progenitor stage. This developmental block was released by iron supplementation (Figure 7E,F). These data are consistent with the requirement for *gata1* for *globin* gene transcription, but also point to the importance of *Mfrn1* for maintenance or expansion of *gata1*^*+*^ erythroid progenitors. The proliferative phenotypes in erythroid progenitor populations occurred during terminal differentiation at about 3 dpf; an analysis of lcr:GFP^+^/gata1:dsRed^+^ and gata1:dsRed^+^ erythroid progenitors at 2 dpf indicated that there were no cell cycle defects at that time point (Supplemental Figure 12).

Lastly, we asked if genetic iron deficiency increased thrombocytosis, akin to the megakaryocytic bias in iron deficient mice ^48^. We crossed *frs* or *weh* mutants with *Tg*(*cd41:GFP*) zebrafish^56^. CD41 is expressed in early hematopoietic progenitors and thrombocytes. 4 dpf *Tg*(*cd41:GFP*) embryos have hematopoietic cells expressing G FP corresponding to hematopoietic progenitor (GFP^lo^) or thrombocyte (GFP^hi^) populations^56^. We did not observe differences in WT vs. *frs* mutants in overall cd41:GFP^+^ (Figure 7G), GFP^lo^ (Figure 7H) or GFP^hi^ (Figure 7I) proportions; further, these percentages were unaltered by iron treatment (gating strategy in Supplemental Figure 13). Similarly, proportions of *cd41*-expressing cells were unaltered in *weh* mutants (Figures 7J-L). Collectively, these data indicate the decrease in erythroid cells in *frs/frs* mutants was not caused by thrombocytic bias. More broadly, they suggest that hematopoietic lineage fate regulation by iron is not conserved between vertebrate species, or that iron is important for adult hematopoietic fate decisions^47,48^ rather than in embryonic hematopoiesis, the context of this study.

Earlier studies with Friend virus infected murine erythroleukemia cell lines recapitulated the hemoglobinization and mitochondrial heme defect, but because these cells have severe defects in cell cycle regulation, we could not rely on those cells to recapitulate these studies in a mammalian system ^18^. We then attempted to recapitulate these studies in a HUDEP2 cells, a human cell line which could recapitulate the requirement for proliferation during differentiation, culminating in quiescence after terminal differentiation^57^. We knocked out *Mfrn1* in these cells; prior to induction of differentiation, we did not observe any cell cycle defects, similar to our observations in the zebrafish (Supplemental Figure 14). Upon induction of differentiation, *Mfrn1*^*-/-*^ cells rapidly underwent apoptosis making it impossible to collect cell cycle data. It is possible that apoptosis might have occurred downstream of G2/M arrest. These observations recapitulate our findings, and previously published data in other tissues, that iron transport via MFRN proteins is essential for cell cycle regulation in actively proliferating primary cells^58^.

## Discussion

Despite the centrality of iron metabolism in erythropoiesis, the regulatory mechanisms remain opaque. To determine how mitochondrial iron transport regulates erythroid development, we developed novel approaches to circumvent major technical limitations. Critically, we showed *in vivo* that mitochondrial iron import by MFRN1 plays a critical role for cell cycle progression past the G2/M checkpoint in terminally differentiating erythroid cells (Figure 7M, Supplemental Figure 15). In contrast, we did not observe cell cycle defects in *weh/weh* (Fpn1-deficient) erythroid cells.

Iron, and iron-containing cofactors are key components in many cell cycle regulators, including DNA polymerases, primases ^33-35,59^, and ribonucleotide reductase^60^. Heme and Fe-S clusters are synthesized in the mitochondria, which explains why defects in mitochondrial iron transport cause severe cell cycle defects (Supplemental Figure 15). Erythroid, and other cell types undergo cell cycle arrest during iron dysregulation or iron deficiency^61-64^, and erythropoiesis has long been coupled with cell cycle regulation ^43,46,65^. In culture, iron deficiency causes cell cycle arrest at the G1/S transition via upregulation of CDK inhibitors p21 and p27 or increased proteosomal degradation of cyclin D1^61,63,64^, but direct links between iron deficiency and cyclin D1 or p21/p27 regulation have not been identified. We speculate that this phenotype was not observed in zebrafish mutants because local iron concentrations in the erythropoietic niche were higher than in published cell culture studies which used iron chelators.

*Mfrn1* deficiency causes embryonic lethality from anemia, and *Mfrn1* ablation in adult hematopoietic tissues caused a 10-fold increase in erythroid progenitors, with a 50% decrease in adult erythrocytes^7^. These data are consistent with our observation that *gata1*-expressing progenitors are overrepresented in the *frs/frs* erythroid compartment. Seguin *et al*. subsequently reported that deletion of *Mfrn1* and *Mfrn2* caused proliferative defects in hepatocytes and bone marrow derived macrophages, decreasing the regenerative capacity of adult livers that had undergone partial hepatectomy^58^. These data underscore the physiological importance of MFRN1/2 and mitochondrial iron transport in cell cycle regulation. However, given the links between iron deficiency and G1/S arrest, and the importance of iron in ribonucleotide reductase, the rate limiting enzyme of the S phase, the G2/M arrest in *frs/frs* (*mfrn1*) erythroid cells was surprising.

How might mitochondrial iron regulate mitosis? Figure 3 indicates that *frs/frs* erythroid cells have highly enlarged nuclei, consistent with the G2/M arrest of most *frs/frs* erythroid cells (Figure 5). We did not observe any binucleate cells, suggesting that the defect occurs prior to telophase. *fthl* (ferritin heavy chain-like) genes were the most highly upregulated in *frs/frs* erythroid cells, suggesting that ferritin transcription is regulated by mitochondrial iron levels, and ferritin is required for maintenance of the erythroid microtubule cytoskeleton^62^. Another top hit in our list of genes upregulated in *Mfrn1* deficiency was *npm1*, a nucleophosmin involved in centrosome duplication and cell proliferation ^66,67^. *actin* was upregulated, adding to a cluster of differentially upregulated cytoskeletal proteins in *frs/frs* erythroid cells (Supplemental Table 3). Our data predict that mitochondrial iron regulates cytoskeletal processes essential for mitosis; clinically, mitochondrial iron deficiency may cause macrocytic phenotypes in erythroid cells.

A key contribution of our work is development of techniques that enable data collection from small numbers of cells. Underscoring the technical accomplishment of our single embryo studies, only 76 erythroid *frs/frs* erythroid cells were sequenced from 10,000 cells loaded for 10X genomics analysis. Our work significantly increases the experimental rigor of zebrafish experiments as we can now distinguish between individual wild-type and heterozygote animals, enabling quantitation of population variability within individual genotypes. Because the external development of zebrafish embryos allows us to address specific questions that are not feasible in mammalian organisms, these methods significantly add to our ability to study development *in vivo*. The iron supplementation experiments, and cell cycle analyses in this work are currently non-feasible in mammals as mutations for iron metabolism genes cause prenatal lethality.

Our results revealed key differences in zebrafish and mammalian erythropoiesis. Primitive mammalian erythroblasts actively proliferate after lineage commitment^68^. During development, globins, heme synthesis and iron acquisition/metabolism genes are transcriptionally upregulated. Concomitantly, morphological changes such as nuclear condensation occur^33-35,59^. While most of these changes have been observed in definitive erythroblasts, they occur in proliferating primitive erythroblasts as well^69^. In contrast, our studies show that zebrafish erythroid cells contain high levels of hemoglobin at early stages of erythropoiesis. Their nuclear size is relatively constant during terminal erythropoiesis while cytoplasmic area increases and elongates (Figure 1). *Mfrn1* mutant erythroid cells have enlarged nuclei, indicating that nuclear condensation is part of the zebrafish erythropoietic program. Our data suggest that nuclear condensation occurs earlier in zebrafish erythropoietic differentiation^29,33-35^.

Iron deficiency contributes to many disease phenotypes in animal models and human patients whose pathophysiology is poorly understood. We have unveiled a specific role for mitochondrial iron in erythropoietic cell cycle regulation and in the process, designed a series of novel experimental techniques that will greatly increase the rigor of zebrafish experiments and expand their applicability to experiments that are intractable in mammalian models.

Conceptually, we reveal the importance of mitochondrial iron transport in cytosolic and nuclear processes including cell cycle regulation and maintenance of erythroid morphology. Unraveling the complexities of intracellular iron trafficking is key for understanding the pathophysiology of iron deficiency and therapeutic mechanisms of iron supplementation.

## Data Availability

scRNAseq data are available on GEO (GEO accession: GSE277034; reviewer token: obmhcqkublexpot).

## Acknowledgements

We thank members of the Yien and Tejero labs for discussion and feedback, especially Jesus Tejero (University of Pittsburgh) for his incisive questions. We especially thank Hugh Hammer (University of Pittsburgh) and his team, and Gwen Talham (University of Delaware) for assistance with zebrafish husbandry, and the late Richard West (University of Delaware) for extensive technical assistance in developing the flow cytometry techniques in this work. We thank Dewayne Falkner, Ailing Liu and Nan Sheng at the University of Pittsburgh Flow Core for technical assistance. We thank the UPMC Children’s Hospital Genomics and Pitt Single Cell cores for assistance with scRNAseq. We are grateful to Robert Lafyatis and Delphine Gomez for help with scRNAseq experimental design. We thank Enrico Novelli and Samit Ghosh at the University of Pittsburgh for their intellectual engagement and feedback during the project, and Diane Ward and Jerry Kaplan (University of Utah) for critical comments on the manuscript.

This work was supported by the Cooley’s Anemia Foundation (Y. Y. Y.) and National Institutes of Health grants R03 DK118307, P01 HL032262, R35 GM1133560 and R01 DK137107 (Y. Y. Y.), R35 GM137976 (A.N.S.), and T32 (Mark Perfetto; PI Novelli—T32 HL0535120).

## References

1. Kassebaum NJ. The Global Burden of Anemia. Hematol Oncol Clin North Am. Apr 2016;30(2):247–308. doi:10.1016/j.hoc.2015.11.002

2. Kassebaum NJ, Jasrasaria R, Naghavi M, et al. A systematic analysis of global anemia burden from 1990 to 2010. Blood. Jan 30 2014;123(5):615–24. doi:10.1182/blood-2013-06-508325

3. Sangkhae V, Fisher AL, Wong S, et al. Effects of maternal iron status on placental and fetal iron homeostasis. J Clin Invest. Feb 3 2020;130(2):625–640. doi:10.1172/jci127341

4. Yien YY, Robledo RF, Schultz IJ, et al. TMEM14C is required for erythroid mitochondrial heme metabolism. J Clin Invest. Oct 2014;124(10):4294–304. doi:10.1172/JCI76979

5. Yien YY, Shi J, Chen C, et al. FAM210B is an erythropoietin target and regulates erythroid heme synthesis by controlling mitochondrial iron import and ferrochelatase activity. J Biol Chem. Dec 21 2018;293(51):19797–19811. doi:10.1074/jbc.RA118.002742

6. Rondelli CM, Perfetto M, Danoff A, et al. The ubiquitous mitochondrial protein unfoldase CLPX regulates erythroid heme synthesis by control of iron utilization and heme synthesis enzyme activation and turnover. J Biol Chem. Aug 2021;297(2):100972. doi:10.1016/j.jbc.2021.100972

7. Troadec MB, Warner D, Wallace J, et al. Targeted deletion of the mouse Mitoferrin1 gene: from anemia to protoporphyria. Blood. May 19 2011;117(20):5494–502. doi:10.1182/blood-2010-11-319483

8. Shaw GC, Cope JJ, Li L, et al. Mitoferrin is essential for erythroid iron assimilation. Nature. Mar 2 2006;440(7080):96–100. doi:10.1038/nature04512

9. Fraenkel PG, Traver D, Donovan A, Zahrieh D, Zon LI. Ferroportin1 is required for normal iron cycling in zebrafish. J Clin Invest. Jun 2005;115(6):1532–41. doi:10.1172/jci23780

10. Brownlie A, Donovan A, Pratt SJ, et al. Positional cloning of the zebrafish sauternes gene: a model for congenital sideroblastic anaemia. Nat Genet. Nov 1998;20(3):244–50. doi:10.1038/3049

11. Brion LP, Heyne R, Lair CS. Role of zinc in neonatal growth and brain growth: review and scoping review. Pediatr Res. May 2021;89(7):1627–1640. doi:10.1038/s41390-020-01181-z

12. Camaschella C, Pagani A, Silvestri L, Nai A. The mutual crosstalk between iron and erythropoiesis. Int J Hematol. Aug 2022;116(2):182–191. doi:10.1007/s12185-022-03384-y

13. Deur CJ, Stone MJ, Frenkel EP. Trace metals in hematopoiesis. Am J Hematol. Nov 1981;11(3):309–31. doi:10.1002/ajh.2830110313

14. Piskin E, Cianciosi D, Gulec S, Tomas M, Capanoglu E. Iron Absorption: Factors, Limitations, and Improvement Methods. ACS Omega. Jun 21 2022;7(24):20441–20456. doi:10.1021/acsomega.2c01833

15. Wang C, Zhang R, Wei X, Lv M, Jiang Z. Metalloimmunology: The metal ion-controlled immunity. Adv Immunol. 2020;145:187–241. doi:10.1016/bs.ai.2019.11.007

16. Wessels I, Fischer HJ, Rink L. Dietary and Physiological Effects of Zinc on the Immune System. Annu Rev Nutr. Oct 11 2021;41:133–175. doi:10.1146/annurev-nutr-122019-120635

17. Shepherd RE, Kreinbrink AC, Njimoh CL, Vali SW, Lindahl PA. Yeast Mitochondria Import Aqueous Fe(II) and, When Activated for Iron-Sulfur Cluster Assembly, Export or Release Low-Molecular-Mass Iron and Also Export Iron That Incorporates into Cytosolic Proteins. J Am Chem Soc. Jun 28 2023;145(25):13556–13569. doi:10.1021/jacs.2c13439

18. Grillo AS, SantaMaria AM, Kafina MD, et al. Restored iron transport by a small molecule promotes absorption and hemoglobinization in animals. Science. May 12 2017;356(6338):608–616. doi:10.1126/science.aah3862

19. Froschauer EM, Schweyen RJ, Wiesenberger G. The yeast mitochondrial carrier proteins Mrs3p/Mrs4p mediate iron transport across the inner mitochondrial membrane. Biochim Biophys Acta. May 2009;1788(5):1044–50. doi:10.1016/j.bbamem.2009.03.004

20. Foury F, Roganti T. Deletion of the mitochondrial carrier genes MRS3 and MRS4 suppresses mitochondrial iron accumulation in a yeast frataxin-deficient strain. J Biol Chem. Jul 5 2002;277(27):24475–83. doi:10.1074/jbc.M111789200

21. Mühlenhoff U, Stadler JA, Richhardt N, et al. A specific role of the yeast mitochondrial carriers MRS3/4p in mitochondrial iron acquisition under iron-limiting conditions. J Biol Chem. Oct 17 2003;278(42):40612–20. doi:10.1074/jbc.M307847200

22. Zhang Y, Lyver ER, Knight SA, Lesuisse E, Dancis A. Frataxin and mitochondrial carrier proteins, Mrs3p and Mrs4p, cooperate in providing iron for heme synthesis. J Biol Chem. May 20 2005;280(20):19794–807. doi:10.1074/jbc.M500397200

23. Donovan A, Brownlie A, Zhou Y, et al. Positional cloning of zebrafish ferroportin1 identifies a conserved vertebrate iron exporter. Nature. Feb 17 2000;403(6771):776–81. doi:10.1038/35001596

24. Donovan A, Lima CA, Pinkus JL, et al. The iron exporter ferroportin/Slc40a1 is essential for iron homeostasis. Cell Metab. Mar 2005;1(3):191–200. doi:10.1016/j.cmet.2005.01.003

25. Butler A, Hoffman P, Smibert P, Papalexi E, Satija R. Integrating single-cell transcriptomic data across different conditions, technologies, and species. Nat Biotechnol. Jun 2018;36(5):411–420. doi:10.1038/nbt.4096

26. Rueb KF, Stachura DL. Using Flow Cytometry to Detect and Quantitate Altered Blood Formation in the Developing Zebrafish. J Vis Exp. Apr 29 2021;(170)doi:10.3791/61035

27. Kulkeaw K, Inoue T, Ishitani T, Nakanishi Y, Zon LI, Sugiyama D. Purification of zebrafish erythrocytes as a means of identifying a novel regulator of haematopoiesis. Br J Haematol. Feb 2018;180(3):420–431. doi:10.1111/bjh.15048

28. Lv P, Liu F. Heme-deficient primitive red blood cells induce HSPC ferroptosis by altering iron homeostasis during zebrafish embryogenesis. Development. Oct 15 2023;150(20)doi:10.1242/dev.201690

29. Ganis JJ, Hsia N, Trompouki E, et al. Zebrafish globin switching occurs in two developmental stages and is controlled by the LCR. Dev Biol. Jun 15 2012;366(2):185–94. doi:10.1016/j.ydbio.2012.03.021

30. Brönnimann D, Annese T, Gorr TA, Djonov V. Splitting of circulating red blood cells as an in vivo mechanism of erythrocyte maturation in developing zebrafish, chick and mouse embryos. J Exp Biol. Aug 10 2018;221(Pt 15)doi:10.1242/jeb.184564

31. Orkin SH, Zon LI. Hematopoiesis: an evolving paradigm for stem cell biology. Cell. Feb 22 2008;132(4):631–44. doi:10.1016/j.cell.2008.01.025

32. Davidson AJ, Zon LI. The ‘definitive’ (and ‘primitive’) guide to zebrafish hematopoiesis. Oncogene. Sep 20 2004;23(43):7233–46. doi:10.1038/sj.onc.1207943

33. An X, Schulz VP, Li J, et al. Global transcriptome analyses of human and murine terminal erythroid differentiation. Blood. May 29 2014;123(22):3466–77. doi:10.1182/blood-2014-01-548305

34. Hu J, Liu J, Xue F, et al. Isolation and functional characterization of human erythroblasts at distinct stages: implications for understanding of normal and disordered erythropoiesis in vivo. Blood. Apr 18 2013;121(16):3246–53. doi:10.1182/blood-2013-01-476390

35. Liu J, Zhang J, Ginzburg Y, et al. Quantitative analysis of murine terminal erythroid differentiation in vivo: novel method to study normal and disordered erythropoiesis. Blood. Feb 21 2013;121(8):e43–9. doi:10.1182/blood-2012-09-456079

36. Petrat F, Weisheit D, Lensen M, de Groot H, Sustmann R, Rauen U. Selective determination of mitochondrial chelatable iron in viable cells with a new fluorescent sensor. Biochem J. Feb 15 2002;362(Pt 1):137-47. doi:10.1042/0264-6021:3620137

37. Chen C, Garcia-Santos D, Ishikawa Y, et al. Snx3 regulates recycling of the transferrin receptor and iron assimilation. Cell Metab. Mar 5 2013;17(3):343–52. doi:10.1016/j.cmet.2013.01.013

38. Ohgami RS, Campagna DR, Greer EL, et al. Identification of a ferrireductase required for efficient transferrin-dependent iron uptake in erythroid cells. Nat Genet. Nov 2005;37(11):1264–9. doi:10.1038/ng1658

39. Zhang DL, Ghosh MC, Ollivierre H, Li Y, Rouault TA. Ferroportin deficiency in erythroid cells causes serum iron deficiency and promotes hemolysis due to oxidative stress. Blood. Nov 8 2018;132(19):2078–2087. doi:10.1182/blood-2018-04-842997

40. Martens K, DeLoughery TG. Sex, lies, and iron deficiency: a call to change ferritin reference ranges. Hematology Am Soc Hematol Educ Program. Dec 8 2023;2023(1):617–621. doi:10.1182/hematology.2023000494

41. Rushton DH, Barth JH. What is the evidence for gender differences in ferritin and haemoglobin? Crit Rev Oncol Hematol. Jan 2010;73(1):1–9. doi:10.1016/j.critrevonc.2009.03.010

42. Gnanapragasam MN, McGrath KE, Catherman S, Xue L, Palis J, Bieker JJ. EKLF/KLF1-regulated cell cycle exit is essential for erythroblast enucleation. Blood. Sep 22 2016;128(12):1631–41. doi:10.1182/blood-2016-03-706671

43. Hwang Y, Futran M, Hidalgo D, et al. Global increase in replication fork speed during a p57(KIP2)-regulated erythroid cell fate switch. Sci Adv. May 2017;3(5):e1700298. doi:10.1126/sciadv.1700298

44. Lu YC, Sanada C, Xavier-Ferrucio J, et al. The Molecular Signature of Megakaryocyte-Erythroid Progenitors Reveals a Role for the Cell Cycle in Fate Specification. Cell Rep. Dec 11 2018;25(11):3229. doi:10.1016/j.celrep.2018.11.075

45. Siatecka M, Lohmann F, Bao S, Bieker JJ. EKLF directly activates the p21WAF1/CIP1 gene by proximal promoter and novel intronic regulatory regions during erythroid differentiation. Mol Cell Biol. Jun 2010;30(11):2811–22. doi:10.1128/mcb.01016-09

46. Socolovsky M. Pas de deux: the coordinated coupling of erythroid differentiation with the cell cycle. Curr Opin Hematol. May 1 2024;31(3):96–103. doi:10.1097/moh.0000000000000811

47. Zhang D, Gao X, Li H, et al. The microbiota regulates hematopoietic stem cell fate decisions by controlling iron availability in bone marrow. Cell Stem Cell. Feb 3 2022;29(2):232-247.e7. doi:10.1016/j.stem.2021.12.009

48. Xavier-Ferrucio J, Scanlon V, Li X, et al. Low iron promotes megakaryocytic commitment of megakaryocytic-erythroid progenitors in humans and mice. Blood. Oct 31 2019;134(18):1547–1557. doi:10.1182/blood.2019002039

49. Ferreira R, Ohneda K, Yamamoto M, Philipsen S. GATA1 function, a paradigm for transcription factors in hematopoiesis. Mol Cell Biol. Feb 2005;25(4):1215–27. doi:10.1128/mcb.25.4.1215-1227.2005

50. Kobayashi M, Yamamoto M. Regulation of GATA1 gene expression. J Biochem. Jul 2007;142(1):1–10. doi:10.1093/jb/mvm122

51. Lyons SE, Lawson ND, Lei L, Bennett PE, Weinstein BM, Liu PP. A nonsense mutation in zebrafish gata1 causes the bloodless phenotype in vlad tepes. Proc Natl Acad Sci U S A. Apr 16 2002;99(8):5454–9. doi:10.1073/pnas.082695299

52. Gutiérrez L, Caballero N, Fernández-Calleja L, Karkoulia E, Strouboulis J. Regulation of GATA1 levels in erythropoiesis. IUBMB Life. Jan 2020;72(1):89–105. doi:10.1002/iub.2192

53. Traver D, Paw BH, Poss KD, Penberthy WT, Lin S, Zon LI. Transplantation and in vivo imaging of multilineage engraftment in zebrafish bloodless mutants. Nat Immunol. Dec 2003;4(12):1238–46. doi:10.1038/ni1007

54. Long Q, Meng A, Wang H, Jessen JR, Farrell MJ, Lin S. GATA-1 expression pattern can be recapitulated in living transgenic zebrafish using GFP reporter gene. Development. Oct 1997;124(20):4105–11. doi:10.1242/dev.124.20.4105

55. Amigo JD, Yu M, Troadec MB, et al. Identification of distal cis-regulatory elements at mouse mitoferrin loci using zebrafish transgenesis. Mol Cell Biol. Apr 2011;31(7):1344–56. doi:10.1128/mcb.01010-10

56. Ma D, Zhang J, Lin HF, Italiano J, Handin RI. The identification and characterization of zebrafish hematopoietic stem cells. Blood. Jul 14 2011;118(2):289–97. doi:10.1182/blood-2010-12-327403

57. Kurita R, Suda N, Sudo K, et al. Establishment of immortalized human erythroid progenitor cell lines able to produce enucleated red blood cells. PLoS One. 2013;8(3):e59890. doi:10.1371/journal.pone.0059890

58. Seguin A, Jia X, Earl AM, et al. The mitochondrial metal transporters mitoferrin1 and mitoferrin2 are required for liver regeneration and cell proliferation in mice. J Biol Chem. Aug 7 2020;295(32):11002–11020. doi:10.1074/jbc.RA120.013229

59. Wong P, Hattangadi SM, Cheng AW, Frampton GM, Young RA, Lodish HF. Gene induction and repression during terminal erythropoiesis are mediated by distinct epigenetic changes. Blood. Oct 20 2011;118(16):e128–38. doi:10.1182/blood-2011-03-341404

60. Ruskoski TB, Boal AK. The periodic table of ribonucleotide reductases. J Biol Chem. Oct 2021;297(4):101137. doi:10.1016/j.jbc.2021.101137

61. Fu D, Richardson DR. Iron chelation and regulation of the cell cycle: 2 mechanisms of posttranscriptional regulation of the universal cyclin-dependent kinase inhibitor p21CIP1/WAF1 by iron depletion. Blood. Jul 15 2007;110(2):752–61. doi:10.1182/blood-2007-03-076737

62. Goldfarb AN, Freeman KC, Sahu RK, et al. Iron control of erythroid microtubule cytoskeleton as a potential target in treatment of iron-restricted anemia. Nat Commun. Mar 12 2021;12(1):1645. doi:10.1038/s41467-021-21938-2

63. Nurtjahja-Tjendraputra E, Fu D, Phang JM, Richardson DR. Iron chelation regulates cyclin D1 expression via the proteasome: a link to iron deficiency-mediated growth suppression. Blood. May 1 2007;109(9):4045–54. doi:10.1182/blood-2006-10-047753

64. Le NT, Richardson DR. Iron chelators with high antiproliferative activity up-regulate the expression of a growth inhibitory and metastasis suppressor gene: a link between iron metabolism and proliferation. Blood. Nov 1 2004;104(9):2967–75. doi:10.1182/blood-2004-05-1866

65. Hidalgo D, Bejder J, Pop R, et al. EpoR stimulates rapid cycling and larger red cells during mouse and human erythropoiesis. Nat Commun. Dec 17 2021;12(1):7334. doi:10.1038/s41467-021-27562-4

66. Okuda M, Horn HF, Tarapore P, et al. Nucleophosmin/B23 is a target of CDK2/cyclin E in centrosome duplication. Cell. Sep 29 2000;103(1):127–40. doi:10.1016/s0092-8674(00)00093-3

67. Tokuyama Y, Horn HF, Kawamura K, Tarapore P, Fukasawa K. Specific phosphorylation of nucleophosmin on Thr(199) by cyclin-dependent kinase 2-cyclin E and its role in centrosome duplication. J Biol Chem. Jun 15 2001;276(24):21529–37. doi:10.1074/jbc.M100014200

68. Vacaru AM, Isern J, Fraser ST, Baron MH. Analysis of primitive erythroid cell proliferation and enucleation using a cyan fluorescent reporter in transgenic mice. Genesis. Nov 2013;51(11):751–62. doi:10.1002/dvg.22420

69. England SJ, McGrath KE, Frame JM, Palis J. Immature erythroblasts with extensive ex vivo self-renewal capacity emerge from the early mammalian fetus. Blood. Mar 3 2011;117(9):2708–17. doi:10.1182/blood-2010-07-299743

