## Supplemental material for "Mitochondrial iron transport via MFRN1 is required for erythroid cell cycle progression"

**Supplemental Methods**

<u>Genotyping</u>

Zebrafish mutant lines were obtained from ZIRC. Genotyping was carried out by PCR amplification of the region flanking the mutation and Sanger sequencing of the PCR product. Primer sequences: *slc25a37*$^{tq223}$ (*frs*): Fwd 5'-CTGCGTGTCTGTGGTAGAAATGTG-3', Rev 5'-CTAACAGCACATTTGACGCATCTC-3'. *slc40a1*$^{tp85c}$ (*weh*): Fwd 5'-AATGGTCATCTCCATTGCTAATATCGC-3', Rev 5'-AAATGTACAATGGCCAAAAAACGAATT-3'.

After the dissociated embryos were passed through the 70uM filter for FACS, embryo debris was washed out with PBS and centrifuged at 17,000 g for 1 minute. The cell pellet was lysed in 20uL of 50mM NaOH at 95°C for 20 minutes and neutralized with 5uL of 1M Tris-HCl (pH 7.4) for PCR analysis.

<u>Staining of sorted erythroid cells</u>

Sorted zebrafish cells were fixed to slides by drying overnight in the dark at RT. The slides were then incubated for 3 minutes with 0.1% Triton X-100 in 1X PBS. The slides were then washed once with 1X PBS. The slides were then incubated with working heme stain solution[1] in the dark for 15 minutes at 37°C in a pre-warmed humidified chamber. The cells were then washed three times in 1X PBS. To stain DNA, the cell were incubated with 2uM Hoescht stain in 1X PBSB in the dark at RT for 15 minutes. The cells were then washed three times in 1X PBS for 3 minutes each and then a final wash in deionized water for 30s. The cells were then mounted with Cytoseal 60 (Epredia 8310-4) and cover slips (Fisher 12544G). The cells were then imaged on an inverted fluorescent stereoscope (Zeiss Axiozoom).

For Wright Giemsa staining, dried slides were incubated with ice-cold methanol for 1 minute and dried overnight in the dark at RT. The slides were stained with 5% Wright-Giemsa for 20 minutes and washed 3 times in de-ionized water for 3 minutes each. The cells were imaged using an inverted stereoscope (Zeiss Axiozoom).

<u>Cell-Cycle Analysis in HUDEP2 cells</u>

A Click-It EdU (ThermoFisher C10419) stock was made following manufacturer's instructions (10mM EdU in DMSO). A final concentration of 10uM Click-It EdU was added to 400,000 HUDEP2 cells/mL. The cells were then cultured under normal conditions for 2 hours (37C, 5% CO2). Cells were then moved onto ice and washed 3X with ice-cold 1X PBS and 1% BSA and centrifuging at 300*g for 5 minutes at 4C. Cells were then fixed with a final concentration of 2% paraformaldehyde. Cells were then washed again 3X with ice-cold 1X PBS and 1% BSA and centrifuging at 300*g for 5 minutes at 4C. Cells were then perforated at room temperature with a final concentration of 0.1% Triton X-100 in 1X PBS with 1% BSA. Cells were then washed 3X with room temperature 1X PBS and 1% BSA and centrifuging at 300*g for 5 minutes at room temperature. The Click-It reaction attaching 647nm Alexa Fluor to EdU following the manufacturer's instruction. Cells were then washed 3X with room temperature 1X PBS and 1% BSA and centrifuging at 300*g for 5 minutes at room temperature. Finally, a final concentration of 1:200 Dye-Cycle Violet (ThermoFisher V35003) was added and passed through a 70um filter. The cells were then analyzed by FACS using the APC and UV (450/50 filter) lasers.

HUDEP2 cells Differentiation and Analysis

Differentiation and analysis were done following the protocols delineated by Vinjamur and Bauer 2018 except for using CD235a-FITC and CD71-APC antibodies (BD Biosciences BDB551374 and BDB559943 respectively) instead of CD235a-APC and CD71-PECy7 antibodies in conjunction with Hoescht33342.

Cas9 RNP-mediated knockout of MFRN1 in HUDEP2 cells via nucleofection

To knock out the *MFRN1* gene in HUDEP2 cells, we used gRNAs designed with Integrated DNA Technologies (IDT) gRNA selection software, prioritizing high specificity for the targeted cut sites. Four gRNAs were selected: three targeting *EXON1* of *hMFRN1* (AA: 5'-ATGACGTTGACGCCTCGCAA, AE: 5'-CTCCGTAGATACTTGTGTAC, AG: 5'-AATGACGTTTTCCACCACCA), and one targeting *EXON2* (AF: 5'-GGCCACCCTGCTCCACGATG). To excise the region spanning EXON1 to EXON2, we paired one of the EXON1-targeting gRNAs (AA, AE, or AG) with the EXON2-targeting gRNA (AF).

Nucleofection was performed as described previously [2]. Briefly, 75 pmol of Cas9 protein was diluted in 3.75 µL of Cas9 buffer (20 mM HEPES pH 7.5, 150 mM KCl, 1 mM $MgCl_2$, 10% glycerol, 1 mM TCEP). A 1.3-fold molar excess of gRNA was prepared in the same buffer, with each gRNA contributing equally to the total when used in pairs. The gRNA solution was added dropwise to the Cas9 buffer over 30 seconds, followed by incubation at room temperature for at least 5 minutes to allow RNP complex formation.

HUDEP2 cells (100,000–200,000) were resuspended in 20 µL of Lonza P3 buffer (Fisher V4XP3032) and mixed with 7.5 µL of the Cas9/gRNA RNP solution. This mixture was transferred into a Lonza S electroporation cuvette (Fisher V4XP3032) and nucleofected using a Lonza 4D Nucleofector with program code DD100. Post-nucleofection, cells were diluted into HUDEP2 complete media at a density of 200,000 to 1,000,000 cells/mL and allowed to recover for at least 12 hours. Cells were then plated into 96-well plates for clonal expansion.

Genotyping was performed using PCR with primers (Forward: 5'-ACACGAATGCAGAGTTTGAGT; Reverse: 5'-CTTCTGCTGGATTCATTACC), and gene knockout was further confirmed by quantitative real-time PCR using probe sets targeting the excised region.

### Supplemental Figures

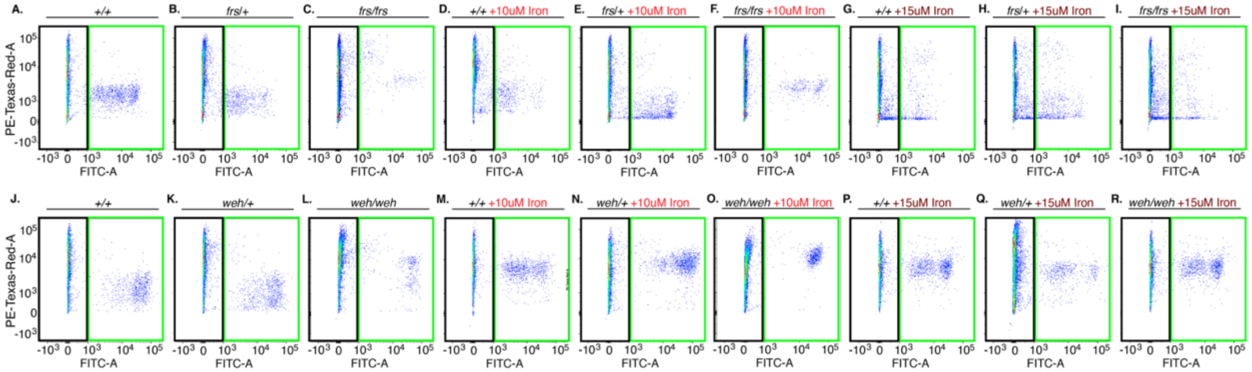


Supplemental Figure 1. Gating strategy for zebrafish RPA staining and representative plots. In each FACS plot, the FITC$^{hi}$ population corresponds to the lcr:GFP$^{+}$ population (gating in Figure 1). The FITC$^{neg/lo}$ population corresponds to the GFP$^{-}$ population. The RPA signal is the average of Texas Red signals within either the GFP$^{+}$ or GFP$^{-}$ populations. Representative plots for the following groups or treatment conditions are shown: (A) +/+; Tg(*lcr:GFP*) (B) *frs/*+; Tg(*lcr:GFP*) (C) *frs/frs*; Tg(*lcr:GFP*) (D) +/+; Tg(*lcr:GFP*) + 10 μM Fe-hinokitiol (E) *frs/*+; Tg(*lcr:GFP*) 10 μM Fe-hinokitiol (F) *frs/frs*; Tg(*lcr:GFP*) 10 μM Fe-hinokitiol (G) +/+; Tg(*lcr:GFP*) 15 μM Fe-hinokitiol (H) *frs/*+; Tg(*lcr:GFP*) 15 μM Fe-hinokitiol (I) *frs/frs*; Tg(*lcr:GFP*) 15 μM Fe-hinokitiol. (J) +/+; Tg(*lcr:GFP*) (K) *weh/*+; Tg(*lcr:GFP*) (L) *weh/weh*; Tg(*lcr:GFP*) (M) +/+; Tg(*lcr:GFP*) + 10 μM Fe-hinokitiol (N) *weh/*+; Tg(*lcr:GFP*) 10 μM Fe-hinokitiol (O) *weh/weh*; Tg(*lcr:GFP*) 10 μM Fe-hinokitiol (P) +/+; Tg(*lcr:GFP*) 15 μM Fe-hinokitiol (Q) *weh/*+; Tg(*lcr:GFP*) 15 μM Fe-hinokitiol (R) *weh/weh*; Tg(*lcr:GFP*) 15 μM Fe-hinokitiol.

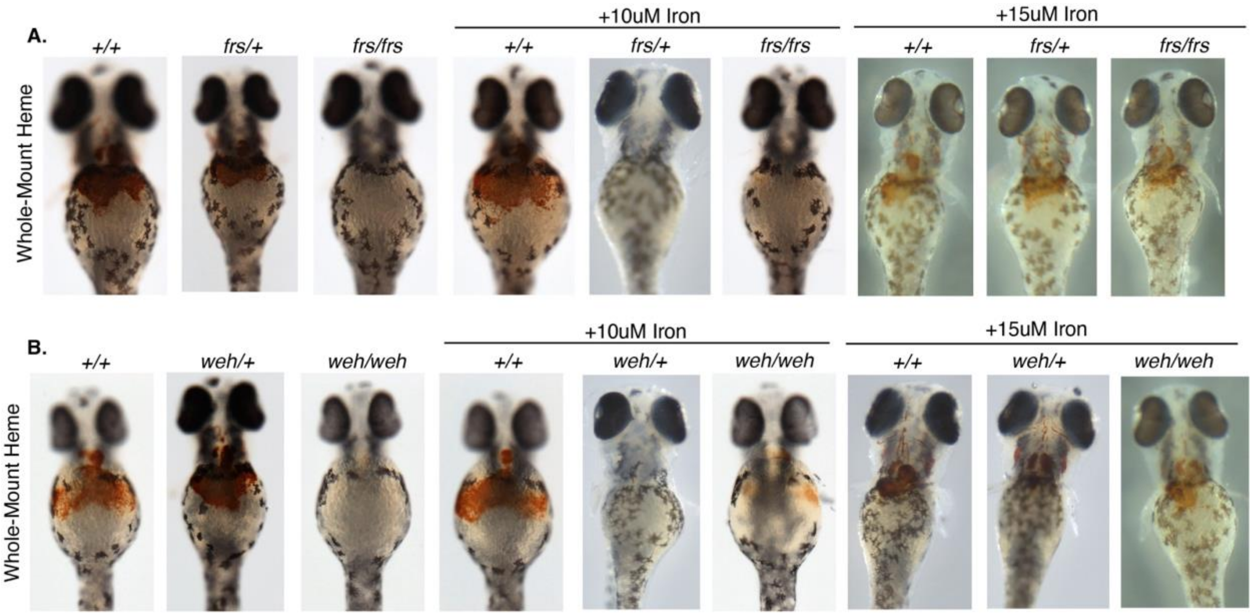


Supplemental Figure 2. Representative benzidine stained images of whole zebrafish obtained from control and Fe-hinokitiol treated groups; genotypes were obtained from incrosses of (A) *frs*/+; Tg(*lcr:GFP*) or (B) *weh*/+; Tg(*lcr:GFP*) zebrafish.

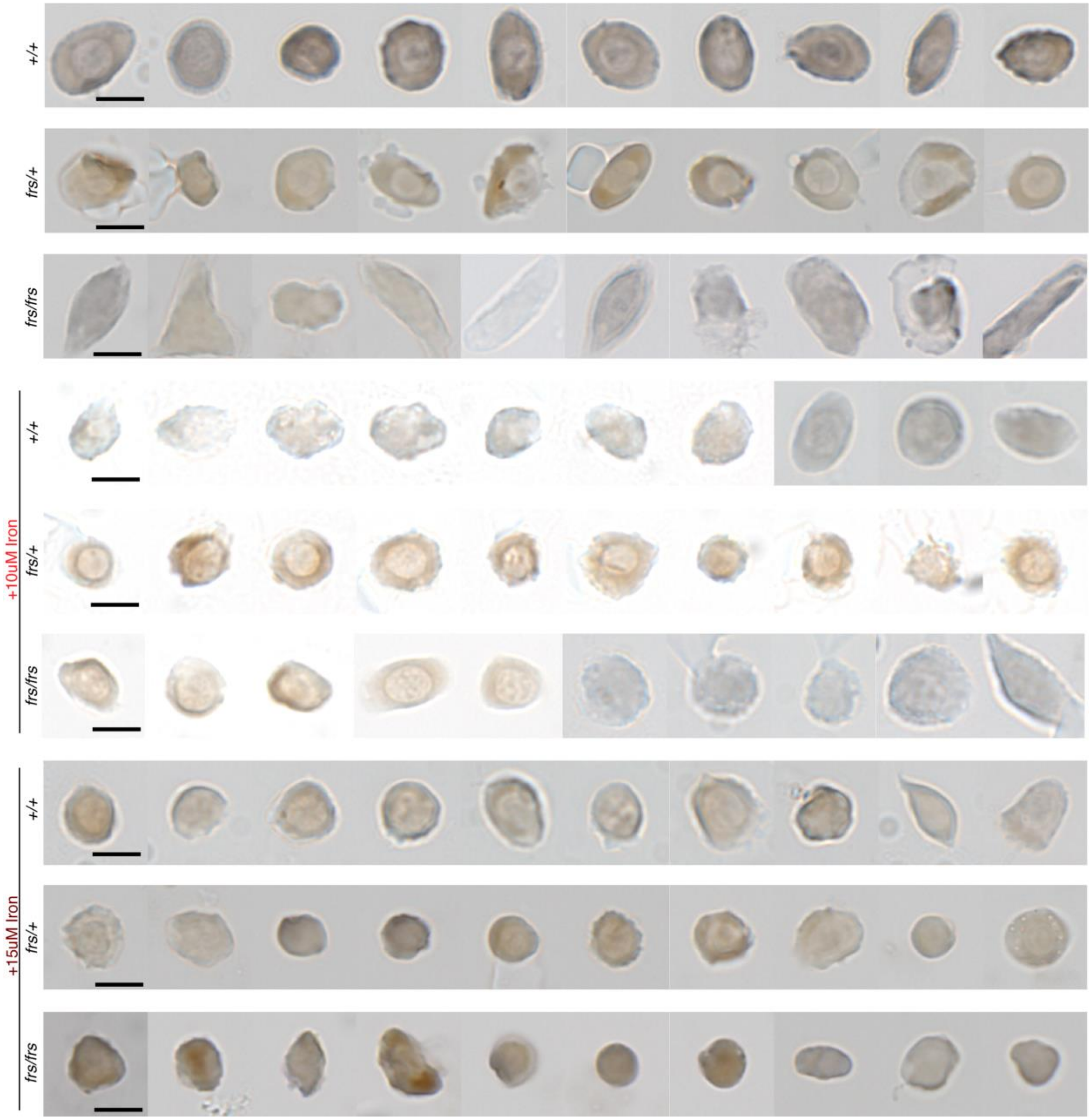


Supplemental Figure 3. Representative benzidine staining of GFP$^+$ erythroid cells from control and Fe-hinokitiol treated WT, *frs/+* and *frs/frs* embryos obtained from incrosses of *frs/+*; Tg(*lcr:GFP*) zebrafish. Scale bar is 5um. Panels consist of cell images spliced together from separate fields because the number of cells on each slide is too small for all the cells to fit onto a page.

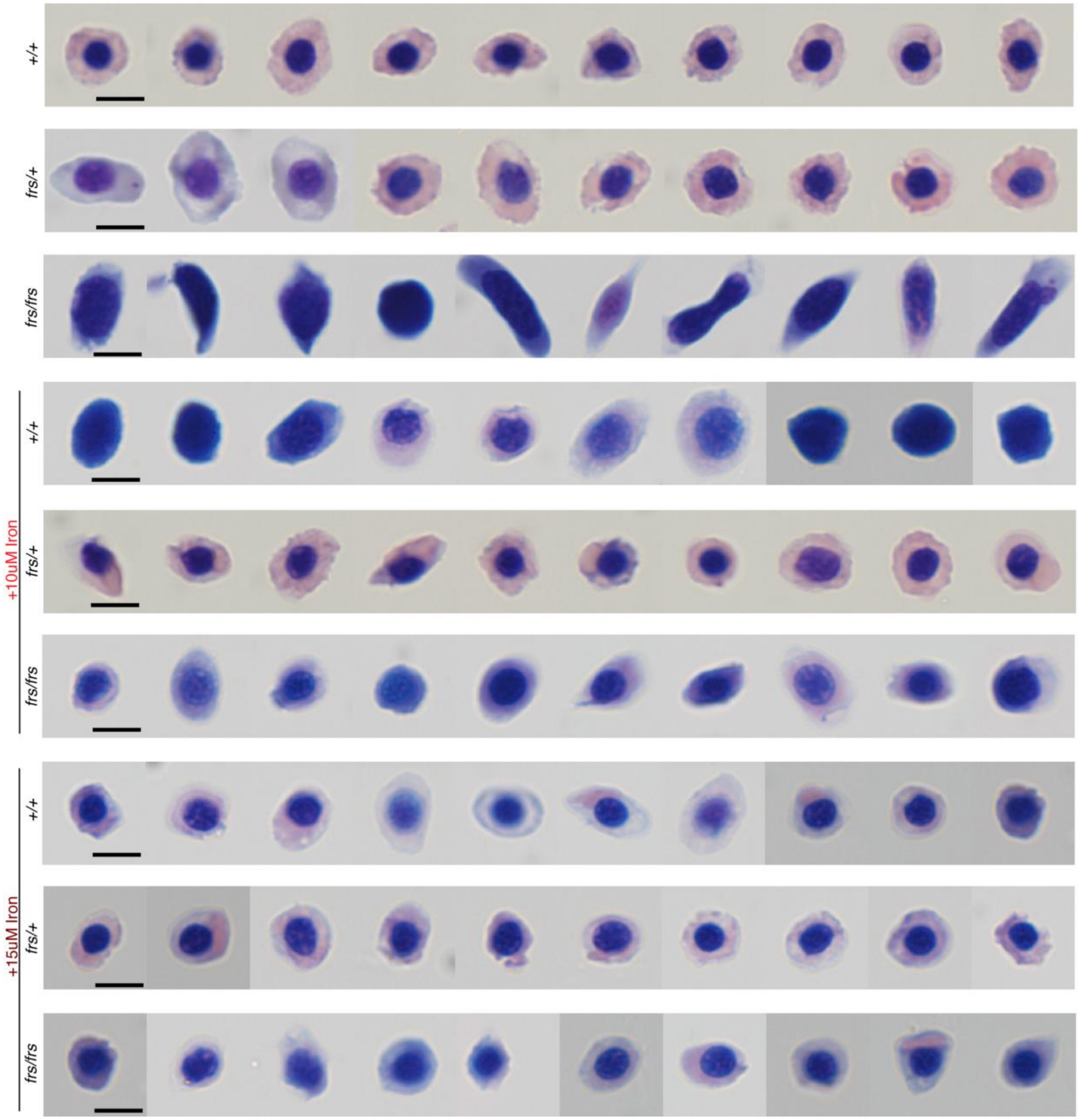


Supplemental Figure 4. Representative Giemsa staining of GFP$^{+}$ erythroid cells from control and Fe-hinokitiol treated WT, *frs/+* and *frs/frs* embryos obtained from incrosses of *frs/+*; Tg(*lcr:GFP*) zebrafish. Scale bar is 5um. Panels consist of cell images spliced together from separate fields because the number of cells on each slide is too small for all the cells to fit onto a page.

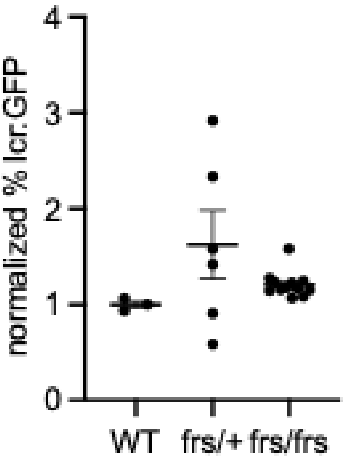


Supplemental Figure 5. *Mfrn1* deficient zebrafish (*frs/frs*) do not have decreases in the number of lcr:GFP expressing cells at 1.5 dpf.

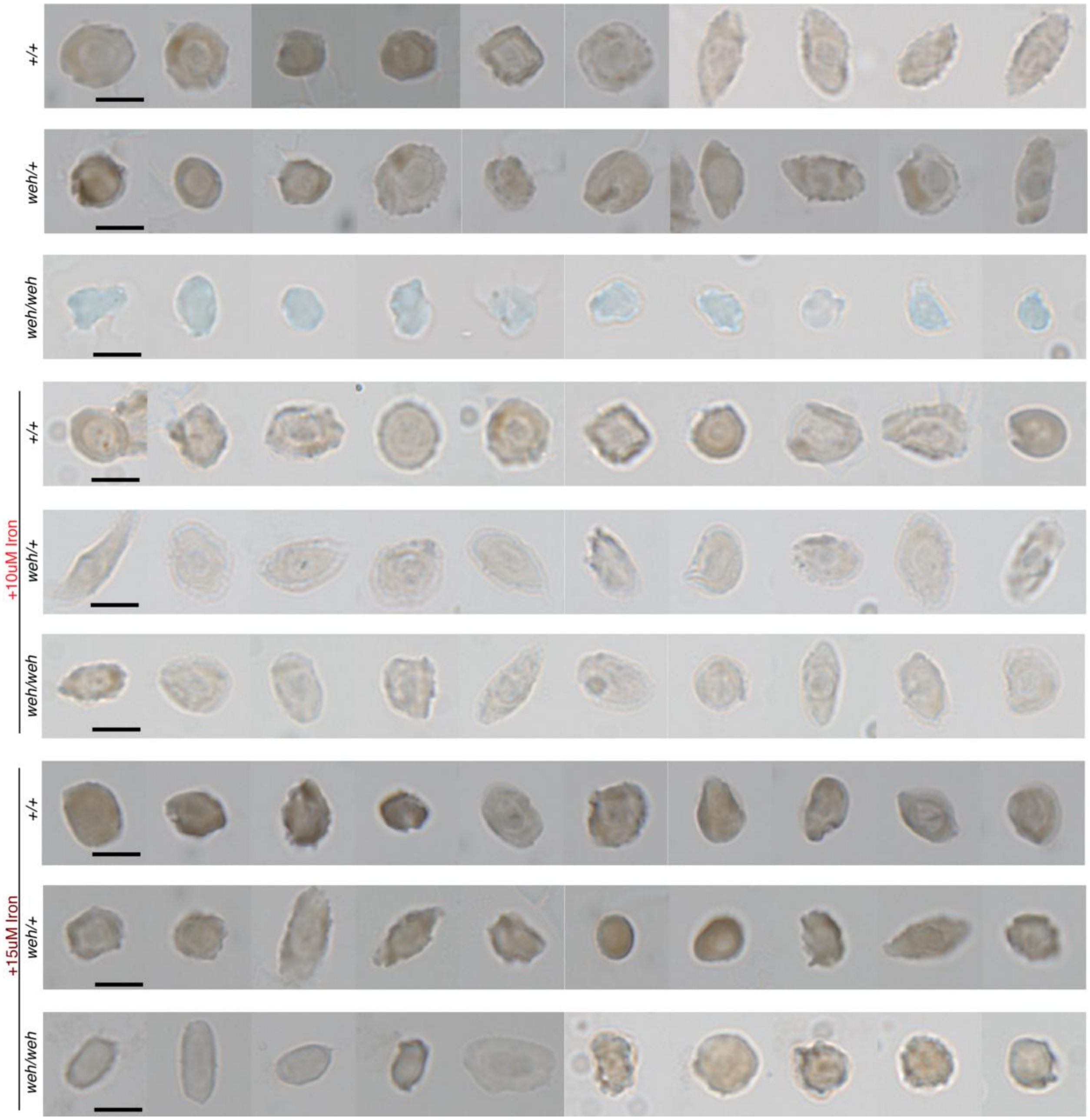


Supplemental Figure 6. Representative benzidine staining of GFP$^+$ erythroid cells from control and Fe-hinokitiol treated WT, *weh*/+ and *weh/weh* embryos obtained from incrosses of *weh*/+; Tg(*lcr:GFP*) zebrafish. Scale bar is 5um. Panels consist of cell images spliced together from separate fields because the number of cells on each slide is too small for all the cells to fit onto a page.

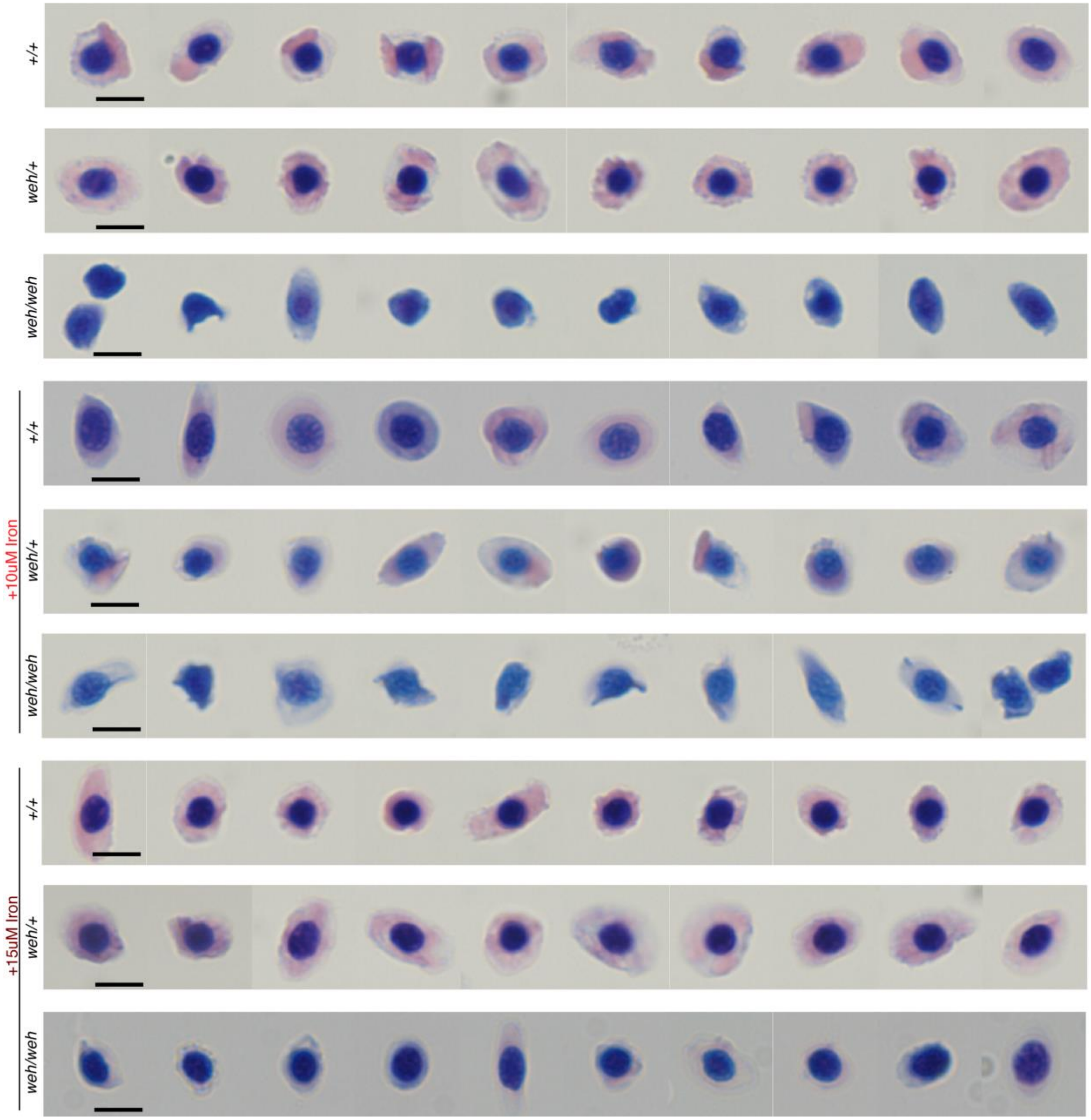


Supplemental Figure 7. Representative Giemsa staining of GFP$^+$ erythroid cells from control and Fe-hinokitiol treated WT, *weh/+* and *weh/weh* embryos obtained from incrosses of *weh/+*; Tg(*lcr:GFP*) zebrafish. Scale bar is 5um. Panels consist of cell images spliced together from separate fields because the number of cells on each slide is too small for all the cells to fit onto a page.

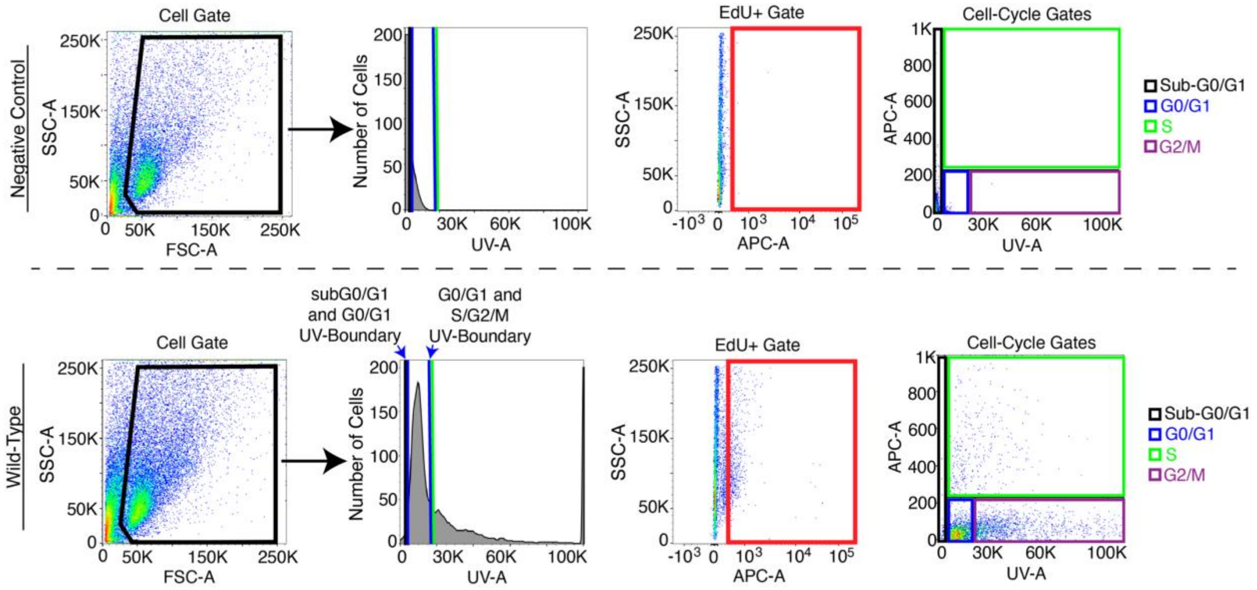


Supplemental Figure 8. Gating strategy for cell cycle analysis of cells obtained from 3 dpf zebrafish embryos. Gating for GFP$^{+}$ and GFP$^{-}$ populations was carried out as described in Figure 1. Analysis of Dyecycle (UV) and EdU+ (APC) gates is shown.

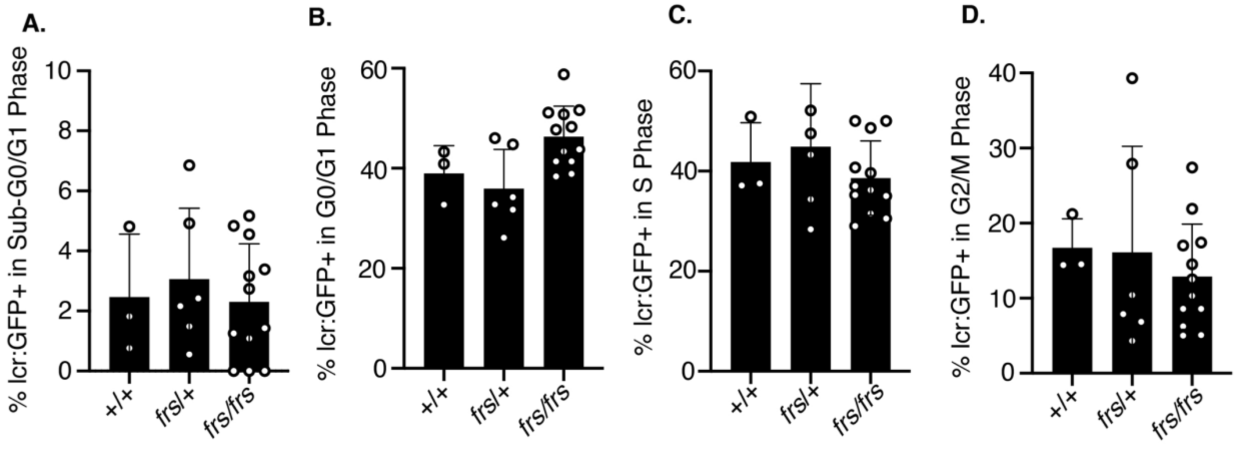


Supplemental Figure 9. At 1.5 dpf, *Mfrn1*-deficient (*frs/frs*) zebrafish embryonic erythroid cells (lcr:GFP expressing) do not have cell cycle defects. (A) apoptosis—sub G0; (B) G0/G1; (C) S phase; (D) G2/M. They are also more proliferative than erythroid cells at 3 dpf.

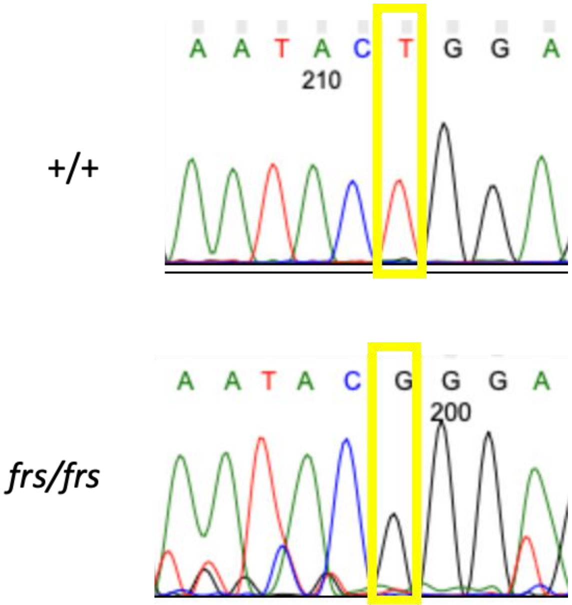


Supplemental Figure 10. Genotyping data obtained from 10, 3-dpf zebrafish embryos pooled together for isolation of cells for scRNA seq analysis.

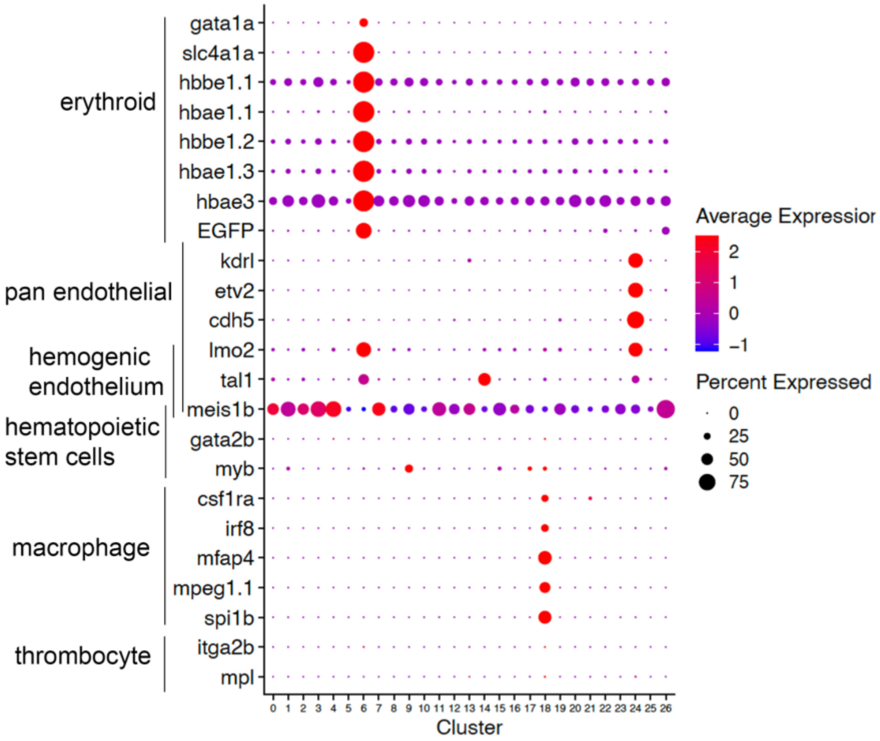


Supplemental Figure 11. Dot plot of gene expression level and frequency of hematopoietic markers in individual clusters.

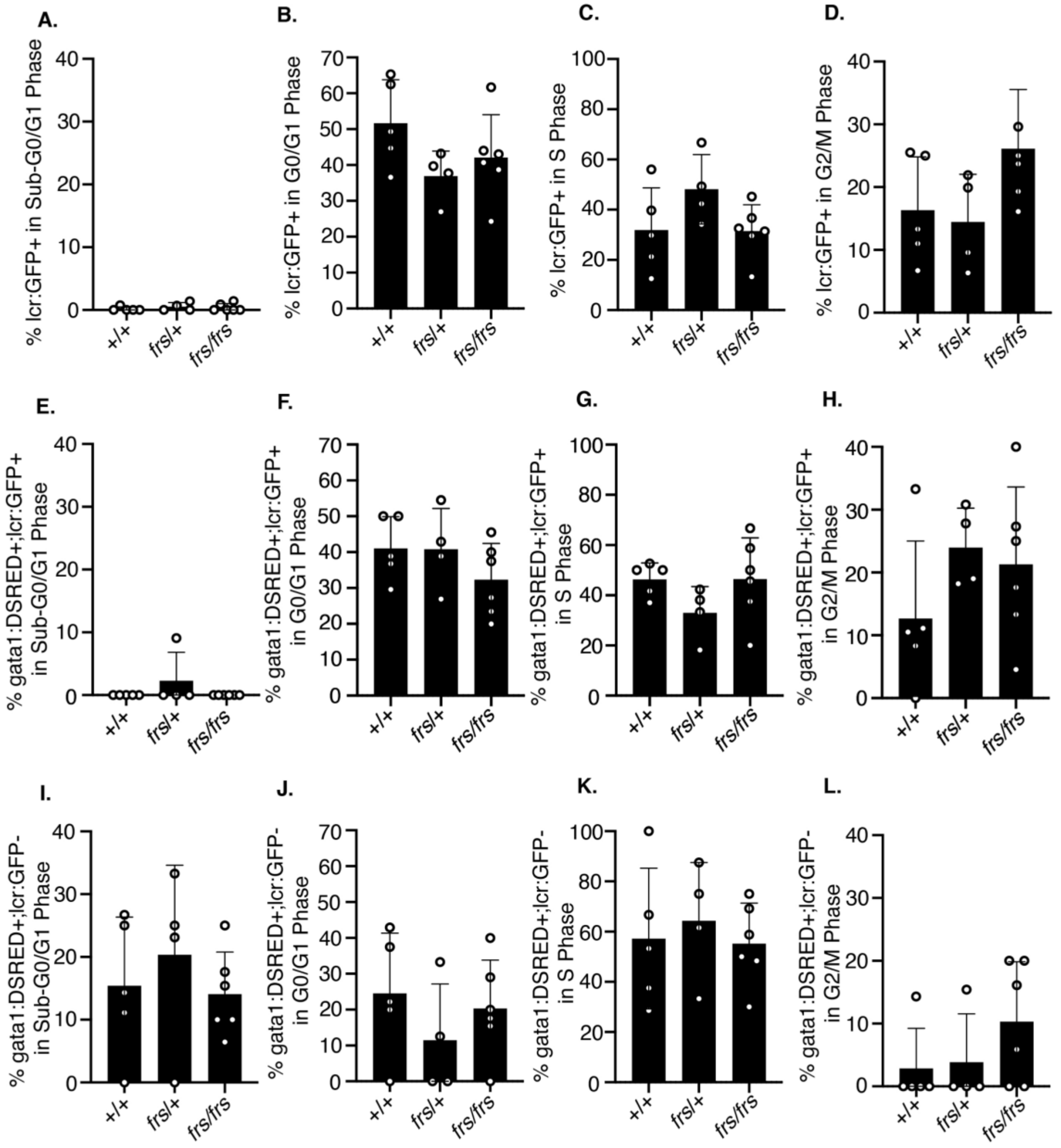


Supplemental Figure 12. At 2 dpf, *Mfrn1*-deficient (*frs/frs*) erythroid cells and progenitors do not exhibit defects in cell cycle regulation relative to wild-type cells. Differentiated cells do not differ in proportions in cell cycle stages and are actively cycling (total lcr:GFP expressing) (A) Sub G0; (B)G1; (C)S phase; (D)G2/M. *Mfrn1*-deficient (*frs/frs*) erythroid progenitors (gata1:dsRed; lcr:GFP expressing) do not exhibit defects in cell cycle regulation relative to wild-type cells and are actively cycling. (E) Sub G0; (F)G1; (G)S phase; (H)G2/M. *Mfrn1*-deficient (*frs/frs*) gata1+ cells that are not yet committed to the erythroid lineage do not exhibit defects in cell cycle regulation relative to wild-type cells. (I) Sub G0; (J)G1; (L)S phase; (L)G2/M.

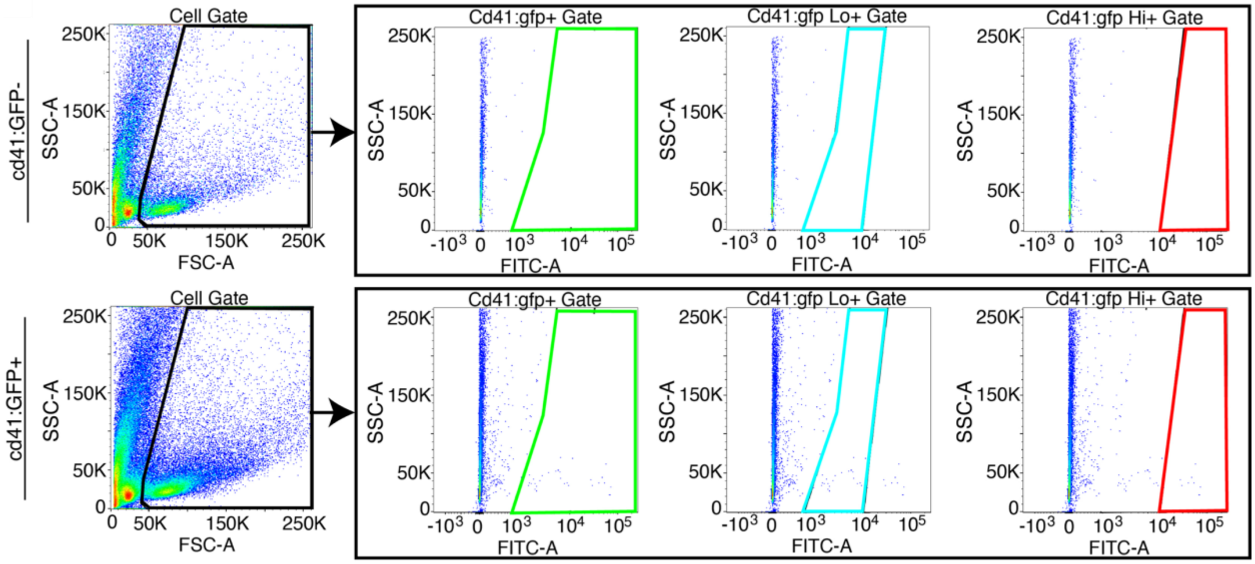


Supplemental Figure 13: Gating scheme for cd41:GFP cell populations. The cd41:GFP$^{+}$ population was defined as the cells not included in the negative control (GFP-). cd41:GFP$^{lo}$ and cd41:GFP$^{hi}$ populations were defined as described [3]

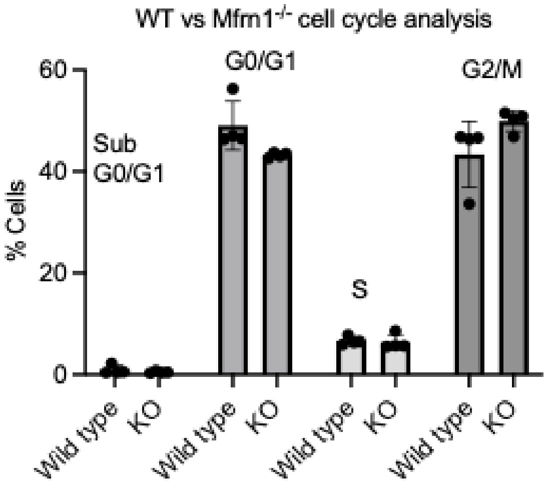


Supplemental Figure 14. Non-differentiating *Mfrn1*$^{-/-}$ HUDEP2 cells do not exhibit cell cycle defects relative to wild-type cells.

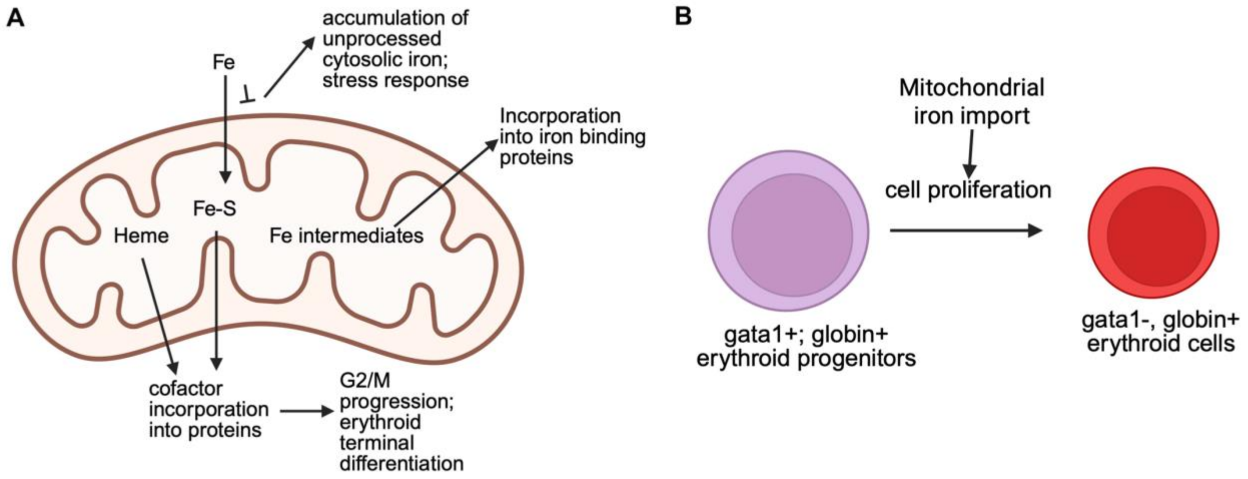


Supplemental Figure 15. Role of mitochondrial iron import in erythropoietic differentiation. (A) Mitochondrial iron is converted into iron intermediates, which are exported into the cytosol and incorporated into iron binding proteins; alternatively, they are used to synthesize Fe-S clusters and heme, which are also exported and incorporated into proteins. This manuscript demonstrates that mitochondrial iron processing is required for G2/M progression in terminally differentiating erythroid cells. When mitochondrial iron transport is impaired, unprocessed iron accumulates in the cytosol, causing stress responses in the cell. (B) Mitochondrial iron import is specifically required for terminal differentiation of gata1$^{+}$; globin$^{+}$ erythroid progenitors into mature gata1$^{-}$; globin$^{+}$ erythroid cells via a process that requires cell proliferation.

### Supplemental References


1. Amigo JD, Ackermann GE, Cope JJ, et al. The role and regulation of friend of GATA-1 (FOG-1) during blood development in the zebrafish. *Blood*. Nov 19 2009;114(21):4654-63. doi:10.1182/blood-2008-12-189910
2. Chung JE, Magis W, Vu J, et al. CRISPR-Cas9 interrogation of a putative fetal globin repressor in human erythroid cells. *PLoS One*. 2019;14(1):e0208237. doi:10.1371/journal.pone.0208237
3. Ma D, Zhang J, Lin HF, Italiano J, Handin RI. The identification and characterization of zebrafish hematopoietic stem cells. *Blood*. Jul 14 2011;118(2):289-97. doi:10.1182/blood-2010-12-327403
